# WNT11 suppresses tumor initiation and invasion by inactivating RAC1

**DOI:** 10.64898/2026.07.13.738348

**Authors:** Subbulakshmi Karthikeyan, Patrick J. Casey, Mei Wang

## Abstract

WNT11, a non-canonical WNT ligand, plays well-defined roles in development and tissue architecture; however, its function in cancer remains ambiguous. Here, we characterize WNT11 as a context-dependent suppressor of cancer stemness, invasion, and *in vivo* tumor formation in human epithelial cancer models. We show that WNT11 upregulation reduces the expression of stemness-promoting genes, suppresses epithelial-to-mesenchymal transition, and inhibits sphere formation and tumor growth. Conversely, downregulation of WNT11 enhances these aggressive malignant properties of cancer cells. Mechanistically, we found that the ability of WNT11 to inhibit RAC1 GTPase activation is essential for its regulation of invasion and self-renewal. In cells unresponsive to WNT11, the connectivity between WNT11 and RAC1 activity is disengaged. Direct manipulation of RAC1 activity in these cells recapitulates the phenotype and molecular signature of WNT11-responsive cells, establishing RAC1 as a critical effector of WNT11-mediated tumor suppression. Taken together, these findings identify the cellular context in which WNT11 suppresses RAC1 activation as a key determinant of its anti-tumor effects and provide a mechanistic framework for understanding the diverse, and sometimes opposing, roles of WNT11 reported in cancer.

## INTRODUCTION

WNT signaling is a highly conserved network essential for embryonic development, stem cell maintenance, and tissue homeostasis. There are 19 WNT ligands and 10 Frizzled receptors that trigger diverse downstream cascades broadly categorized into canonical (β-catenin-dependent) and non-canonical (β-catenin-independent) pathways [1]. While the canonical pathway stabilizes β-catenin and drives transcription of oncogenic target genes to promote stemness [2,3], the non-canonical signaling is known to regulate planar cell polarity (PCP) and WNT/Ca²⁺ pathways that affect cytoskeletal organization, migration, and polarity [4,5]. Dysregulation of both pathways has been implicated in a wide range of cancers, including colorectal, breast, prostate, and liver malignancies [2,6–8].

The development of WNT targeting therapies has largely focused on canonical WNT signaling. However, non-canonical WNT ligand-driven pathways are emerging as critical regulators of tumor plasticity and progression. Among these ligands, WNT11 plays a well-established role in embryogenesis, where it controls convergent extension and polarity through PCP and calcium-dependent mechanisms [9]. However, its role in cancer remains controversial and context dependent. Some studies suggest that WNT11 promotes epithelial-mesenchymal transition (EMT), invasion, and metastasis in cancers [10,11], while others report WNT11 has tumor-suppressive functions, including suppression of β-catenin signaling, promotion of differentiation, and inhibition of tumor growth [12–14]. The diverging roles of WNT11, likely governed by cell signaling context, need to be comprehensively evaluated.

Recent work from our laboratory uncovered a novel connection between WNT11 and RAB4A GTPase. We found that suppressing RAB4A reduces cancer stemness and tumor initiation, and attenuates EMT phenotypes across multiple cancer models [15,16]. Transcriptomic profiling of RAB4A depleted cells identified WNT11 as one of the top upregulated targets [16]. Given the known convergence of non-canonical WNT signaling on Rho family GTPases, including our recent finding of RAC1 being a critical effector of RAB4A driven stemness and invasion, we hypothesized that WNT11 might mediate tumor-suppressive functions by modulating RAC1 activity [14–16].

Multiple inhibitors targeting WNT signaling are in therapeutic development; these agents broadly suppress WNT signaling including both canonical and non-canonical pathways [17–20]. Hence, there is a pressing need for mechanistic understanding of the functional roles of non-canonical WNT ligand signaling in cancers, especially with regard to the cellular context. In the present study, we evaluated the role of WNT11 across a panel of human cancer cell lines. By using both gain- and loss-of-function models, we assessed how WNT11 regulates EMT, invasion, cancer stemness, and tumor formation. Importantly, we have determined that WNT11 functions as a strong tumor suppressor only when its signaling is coupled to RAC1 inactivation.

## RESULTS

### WNT11 is an independent regulator of the cancer stemness program that is silenced in many invasive cancer cell lines

In our recent study, we identified RAB4A as a key regulator of EMT and stemness in several aggressive cancers [16]. Interestingly, WNT11 – a non-canonical pathway WNT ligand – was consistently upregulated when RAB4A is knocked down (Supplementary Fig. S1A), which suggests a potential causal role of WNT11 in suppressing stemness and EMT in cancer [16].

Since WNT11, as other WNT ligands, is a secreted signaling protein, we next extended the evaluation of level of transcription to amount of protein, both in the cell and in the media. To this end, we used the same MDA-MB-231 control and stable RAB4A knockdown isogenic cells as those used for transcription analysis in Supplementary Fig. S1A. Western blotting of the concentrated media revealed a prominent WNT11 band at ∼55 kDa visible only in RAB4A knockdown cells (Supplementary Fig. S1B, group 1) that shifted to the expected intracellular size of ∼39 kDa upon treatment with a deglycosylation enzyme mix [21–23] (Supplementary Fig. S1B, group 2 and 3), thus visualizing the level of post-translationally modified, secreted WNT11. WNT11 levels in whole cell lysates were compared with that in the media (Supplementary Fig. S1B, group 3). The data reveals that knockdown RAB4A upregulates WNT11 production and that the secreted WNT11 level correlates with that in the cells, suggesting that RAB4A regulates WNT11 production rather than secretion.

Since RAB4A has been shown in our earlier study as a strong positive regulator for cancer stemness and EMT, the robust negative correlation of RAB4A and WNT11 suggests a potential opposing role of WNT11 in the regulation in the same process. To set the foundation for the WNT11 function study, we first screened a panel of human epithelial cancer cell lines for endogenous WNT11 expression to identify cell lines for loss and gain of function evaluation. Notably, WNT11 is low or undetectable in aggressive breast (MDA-MB-231), prostate (PC3), glioblastoma (SNB19), and ovarian (OVCAR5) cancer cells. In contrast, higher expression is observed in luminal breast (MCF7), non-small cell lung (H1299), and glioblastoma (U87) cells (Supplementary Fig. S1C). Based on the expression patterns, we generated stable WNT11 overexpressing (OE) lines in WNT11-low cells (Supplementary Fig. S1D), and stable WNT11 knockdown (KD) in WNT11-high cells (Supplementary Fig. S1E). These cells were used for subsequent WNT11-driven function analyses.

To investigate the role of WNT11 in regulating stemness in cancer, we first performed a multiplex qPCR array of a group of experimentally validated Cancer Stem Cell (CSC) related genes (SAB target list H96, Bio-Rad) on the established epithelial cancer cell line pairs with stable modulation of WNT11 expression (Supplementary Fig. S1D–E). In the WNT11-low MDA-MB-231, PC3, and SNB19 cells, enforced WNT11 expression led to striking and broad downregulation of cancer stemness genes such as ALDH1A2, ZEB1, SOX2, OCT4, NANOG, KLF4, EPCAM and BMI1 (Fig. 1A–B; Supplementary Fig. S2A-B). Conversely, silencing WNT11 in WNT11-high MCF7 and H1299 cells resulted in elevated expression of these stemness genes (Fig. 1C–D; Supplementary Fig. S2C-D). These data indicate that WNT11 is a negative regulator for cancer stemness program in several epithelial cancers of different tissue origins.

**Figure 1.**
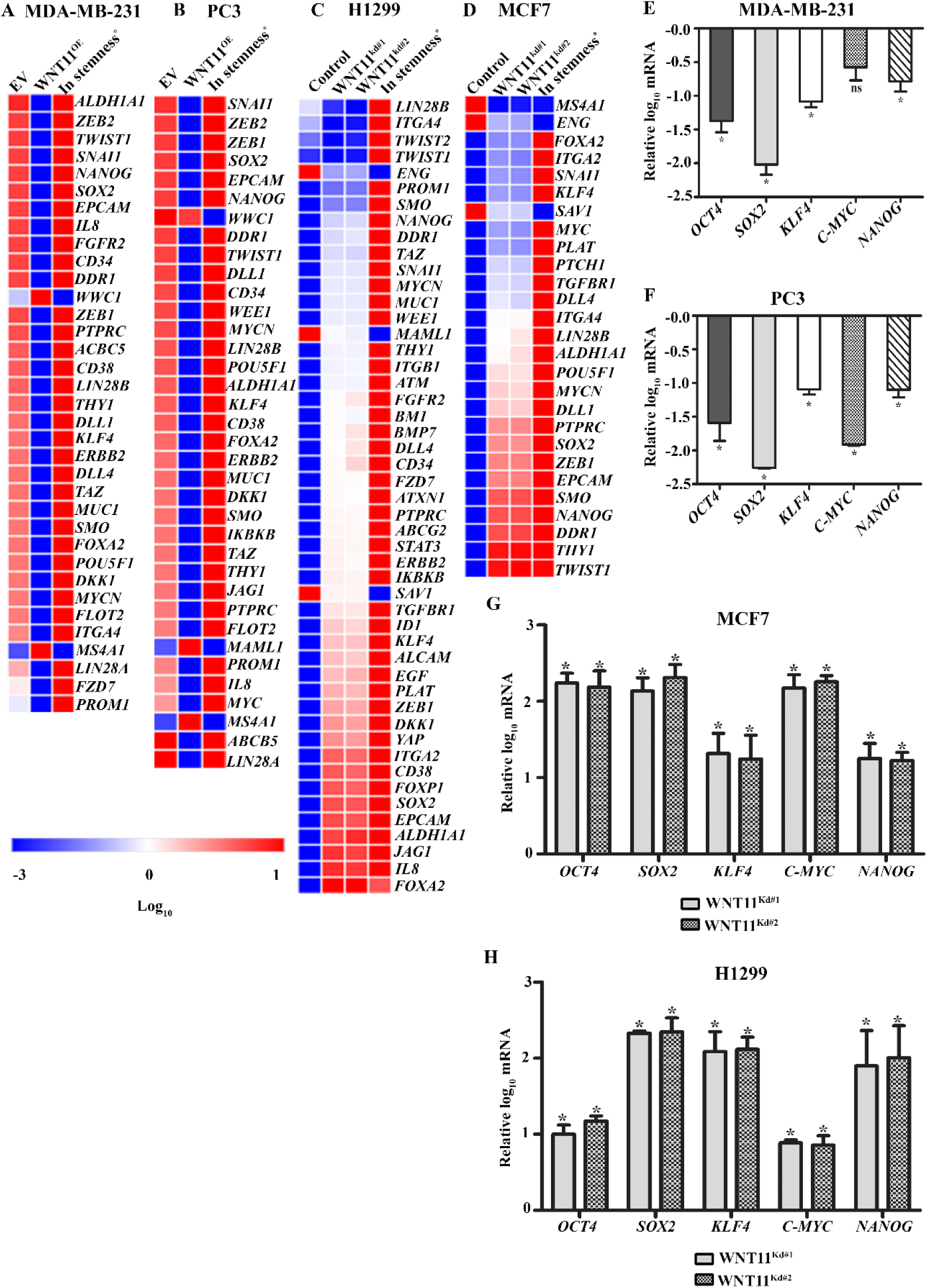
Multiplex gene expression profiling identifies WNT11 as a negative regulator of cancer stemness. **(A-D)** qPCR profiling was performed using a PrimePCR multiplex array comprising 87 experimentally validated genes essential for cancer stemness (SAB target list H96, Bio-Rad). For each cell line that was engineered to either express WNT11 or WNT11-shRNA, relative mRNA expression levels were calculated with respect to control cells (parental cells with empty vector). Genes that were consistently altered in response to WNT11 overexpression in WNT11-low MDA-MB-231 and PC3 cells **(A, B)** or knockdown in MCF7 and H1299 cells **(C, D)** are presented as heat maps, generated using Morpheus software. The last column in each panel indicates the direction of regulation (up or down) in relation to known cancer stemness phenotypes based on published literature [16]. **(E-H)** qPCR analysis for the expression of Yamanaka Factors and NANOG in response to WNT11 overexpression **(E, F)** or knockdown **(G, H)**. Data are presented as mean ± SEM (n ≥ 3; n is the number of biological repeats). *p < 0.05 compared to control cells, which are set as the baseline.

To validate WNT11 impact on the core stemness regulators, we next quantified the expression of the so-called Yamanaka factors (SOX2, OCT4, C-MYC, KLF4), and a closely related stem factor NANOG, by qPCR. Strikingly, WNT11 overexpression in WNT11-low cells led to marked suppression of these genes (Fig. 1E-F; Supplementary Fig. S2E), while WNT11 knockdown in WNT11-high cells significantly increased their expression (Fig. 1G-H). Taken together, these results provide compelling evidence that WNT11 acts as a potent suppressor of stemness programs in epithelial cancer cells across diverse tissue origins. Notably, these transcriptional shifts appeared not to affect short-term proliferation or cell death rates (Supplementary Fig. S3 and Supplementary Fig. S4), consistent with the notion that WNT11 selectively regulates transcriptional programs related to plasticity and stemness. Hence, we next examined the phenotypic impact of WNT11 on stemness-related cancer cell properties.

### WNT11 suppresses cancer cell invasion, EMT, and self-renewal capacity *in vitro*

To assess the functional impact of WNT11 on cancer cell behavior, we performed both invasion and serial replating sphere formation assays using the stable WNT11 overexpression and knockdown cancer cell lines in comparison to the parental cells. Enforced WNT11 expression in WNT11-low cells led to a significant reduction in invasive capacity (Fig. 2A). qPCR analysis revealed a robust downregulation of the mesenchymal marker ZEB1 and a concurrent upregulation of the epithelial marker CDH1 (E-cadherin) (Fig. 2B), consistent with a mesenchymal-to-epithelial shift and reduced invasive potential.

**Figure 2.**
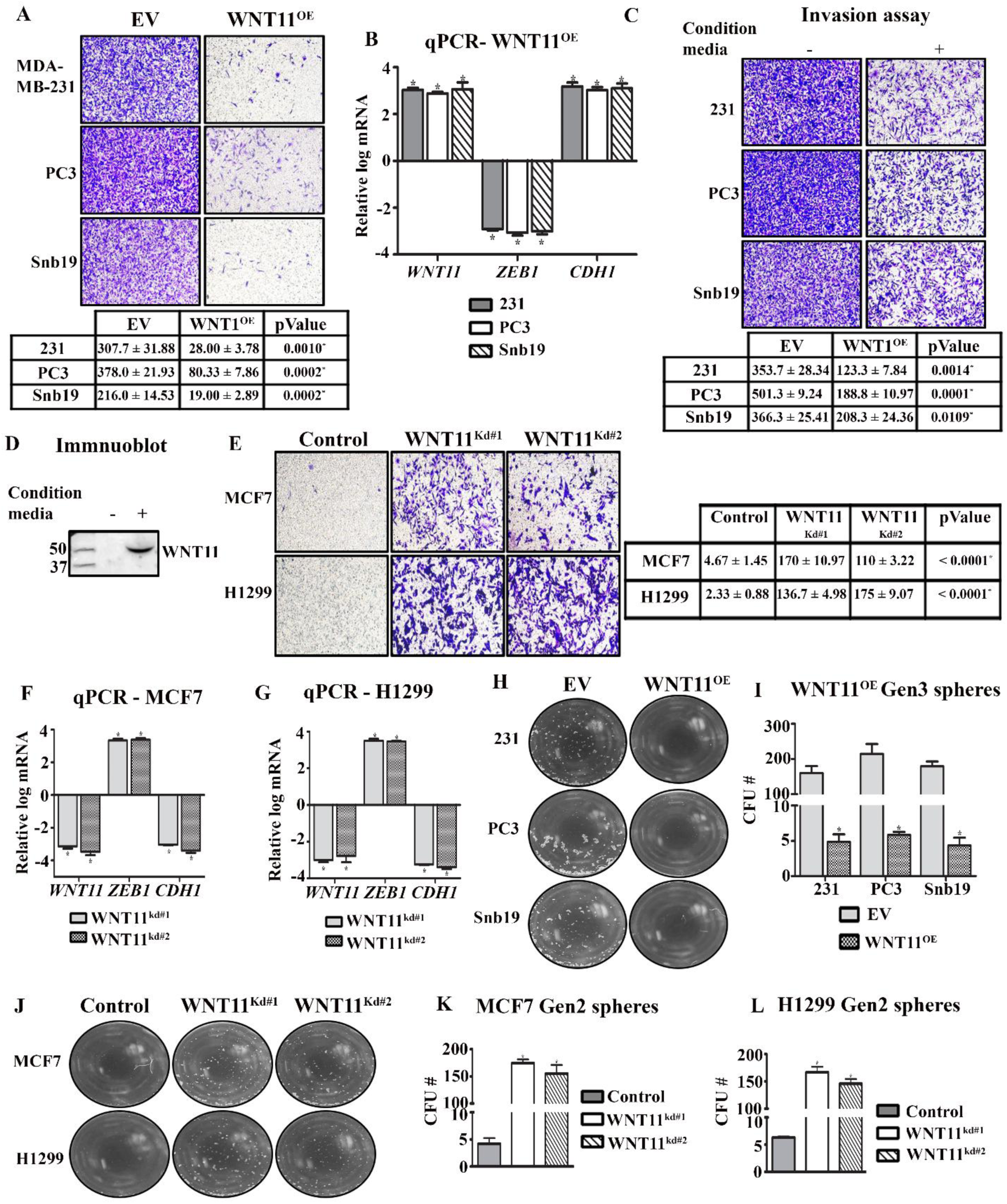
WNT11 levels impact invasive capacity, key EMT gene expression and stemness properties of cancer cells. In all the panels, EV refers to Empty Vector control for exogenous expression. **(A)** Representative images of Matrigel invasion assays performed on WNT11-low cell lines with and without WNT11 overexpression. In all invasion assays, cells were allowed to invade for 24 h prior to visualization. Quantification of invading cells is shown below the images and presented as mean ± SEM, with corresponding p values indicating statistical significance. **(B)** Quantitative PCR validation of key epithelial-mesenchymal transition (EMT) markers in the same cells as in **(A)**; baseline expression is defined by the EV control cells which is set at 0. **(C)** Matrigel invasion assays on the parental WNT11-low cell lines performed in the medium conditioned by WNT11 over-expressing MDA-MB-231 cells. Quantification of invading cells is shown below the images and presented as mean ± SEM, with corresponding p values indicating statistical significance. **(D)** Immunoblot analysis of WNT11 present in the media conditioned by MDA-MB-231 cells with and without stable exogenous expression of WNT11. **(E)** Representative images of Matrigel invasion assays performed on WNT11-high cell lines with or without WNT11-shRNA expression. Quantification of invading cells is shown below the images and presented as mean ± SEM, with corresponding p values indicating statistical significance. **(F, G)** Quantitative PCR validation of key epithelial-mesenchymal transition (EMT) markers in the same cells as in **(E)**. (**H, I**) Serial replating sphere formation studies on WNT11-low cells with or without WNT11 overexpression. **(H)** Representative microscopic images from the assays. **(I)** Quantification of third-generation (Gen-3) spheres in the same cells as in **(G)**. (**J-L**) Serial replating sphere formation studies on WNT11-high cells with or without WNT11 knockdown. **(J)** Representative images of the second-generation (Gen-2) spheres. **(K, L)** Quantification of the spheres in **(I)**. Data are presented as mean ± SEM (n ≥ 3, n is the number of biological replicates). *p < 0.05 compared to control cells, which are set as the baseline.

To validate that it is indeed the secreted WNT11 ligand that is accountable for the inhibitory effect on the invasion, we performed the invasion study using the medium conditioned by WNT11 over-expressing cells. Specifically, media from plates harboring MDA-MB-231 WNT11 over-expressing cells was used as the suspension/top chamber medium in the invasion study on parental MDA-MB-231, PC3 and SNB19 cells. As expected, over-expression of WNT11 in MDA-MB-231 cells elevated its level in the medium compared to that culturing the control cells (Fig. 2D). Consistent with the result from WNT11 over-expressing cells (Fig. 2A), the conditioned medium similarly inhibited the invasion of these highly invasive WNT11-low cells (Fig. 2C). We speculate that the weaker effect of the conditioned medium would be strengthened by concentrating the media.

In contrast to the over-expression study, WNT11knockdown in WNT11-high MCF7 and H1299 cells led to a significant increase in invasion (Fig. 2E). This increase in invasiveness was accompanied by upregulation of ZEB1 and downregulation of CDH1 (Fig. 2F-G), consistent with the conclusion from over-expression study that WNT11 is a negative regulator of the EMT program and cell invasiveness.

To study the effect of WNT11 on cancer stemness, we employed the serial replating sphere formation assay, an established method to evaluate self-renewal capacity. In this assay, only cells with stemness capacity are capable of generating spheres through multiple passages in non-adherent, serum-free conditions. Exogenous WNT11 expression in WNT11-low cells led to a marked decrease in third generation sphere formation (Fig. 2H-I), indicating impaired self-renewal ability. Conversely, WNT11 knockdown in WNT11-high cells resulted in increased secondary sphere formation, consistent with enhanced stem-like traits (Fig. 2J-L).

Taken together, these findings demonstrate that WNT11 plays a tumor suppressive role for EMT, invasion, and self-renewal, across diverse cancer types.

### WNT11 suppresses *in vivo* tumor formation

Having observed the impact of WNT11 modulation on stemness and invasion *in vitro*, we extended the study to *in vivo* tumor formation. To this end, we implanted both WNT11 overexpressing and WNT11 knockdown cells and the respective parental cells subcutaneously in NOD-SCID mice to observe the course of tumor formation. For WNT11-low MDA-MB-231 and PC3 lines, the control cells reliably formed tumors within 3–4 weeks, while their WNT11-overexpressing counterparts failed to grow any detectable tumors after extended observation (until euthanization at 90-day post-implantation) (Fig. 3A, top and bottom panels). In contrast, for the WNT11-high MCF7 and H1299 lines, the control cells, as expected for their indolent nature as previously reported [24–27], formed no visible tumors up to 90 days, while their counterpart with stable knockdown of WNT11 formed tumors by 4–5 weeks (Fig. 3B, top and bottom panels). Prior to implantation, all cell preparations were confirmed to be >95% viable by trypan blue exclusion counting, excluding the possibility that lack of tumor formation was due to poor cell viability. Hence, under both over-expression and knockdown settings of WNT11, we observed striking changes in tumor formation that support the role of WNT11 as an independent tumor suppressor, consistent with the in vitro study results. These observations also highlight the potential of targeting WNT11 specific signaling in cancer treatment.

**Figure 3.**
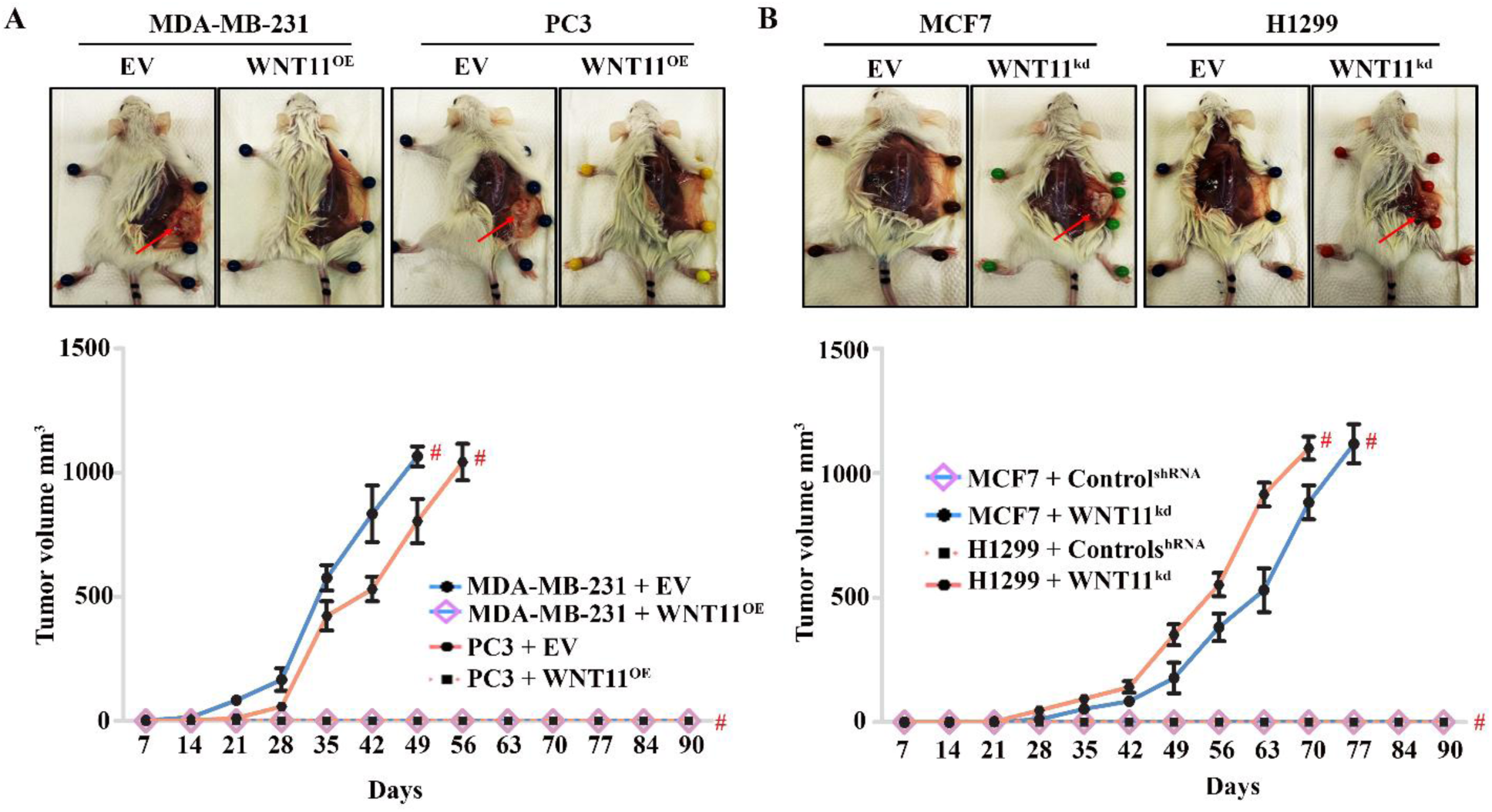
Xenograft tumor formation study. (**A, B**) Top panels display the photographs of representative mice and the tumors (red arrows) at the time of sacrifice. Five mice were implanted with each cell line (N = 5) subcutaneously in the flank region. Bottom panels present the growth curves of the xenograft tumors. The group identities are provided below the graphs. “#” indicates the time point when mice were sacrificed in accordance with IACUC protocol, either upon reaching 1 cm^3^, or at the end of the experiment. EV, Empty Vector control.

### WNT11’s tumor suppression function depends on its ability to inhibit RAC1 activation

Among the cell lines we surveyed for WNT11 expression (Supplementary Fig. S1B), we found exceptions for WNT11 acting as tumor suppressor. In WNT11-low OVCAR5 cells, WNT11 overexpression had no impact on either cell invasion (Fig. 4A) or on the expression of the EMT regulatory genes ZEB1 and CDH1 (Fig. 4B). Similarly, WNT11 knockdown in WNT11-high U87 cells failed to enhance invasion or alter EMT gene expression (Fig. 4C-D). We also looked at the impact of WNT11 on self-renewal ability of the two cell lines by performing serial replating sphere assays. In contrast to the responsive cell lines, WNT11 overexpression in OVCAR5 and its knockdown in U87 cells had no impact on third generation sphere formation (Fig. 4E-F). These findings indicate that OVCAR5 and U87 cells are phenotypically unresponsive to WNT11 modulation, in contrast to the other cell lines studied above.

**Figure 4.**
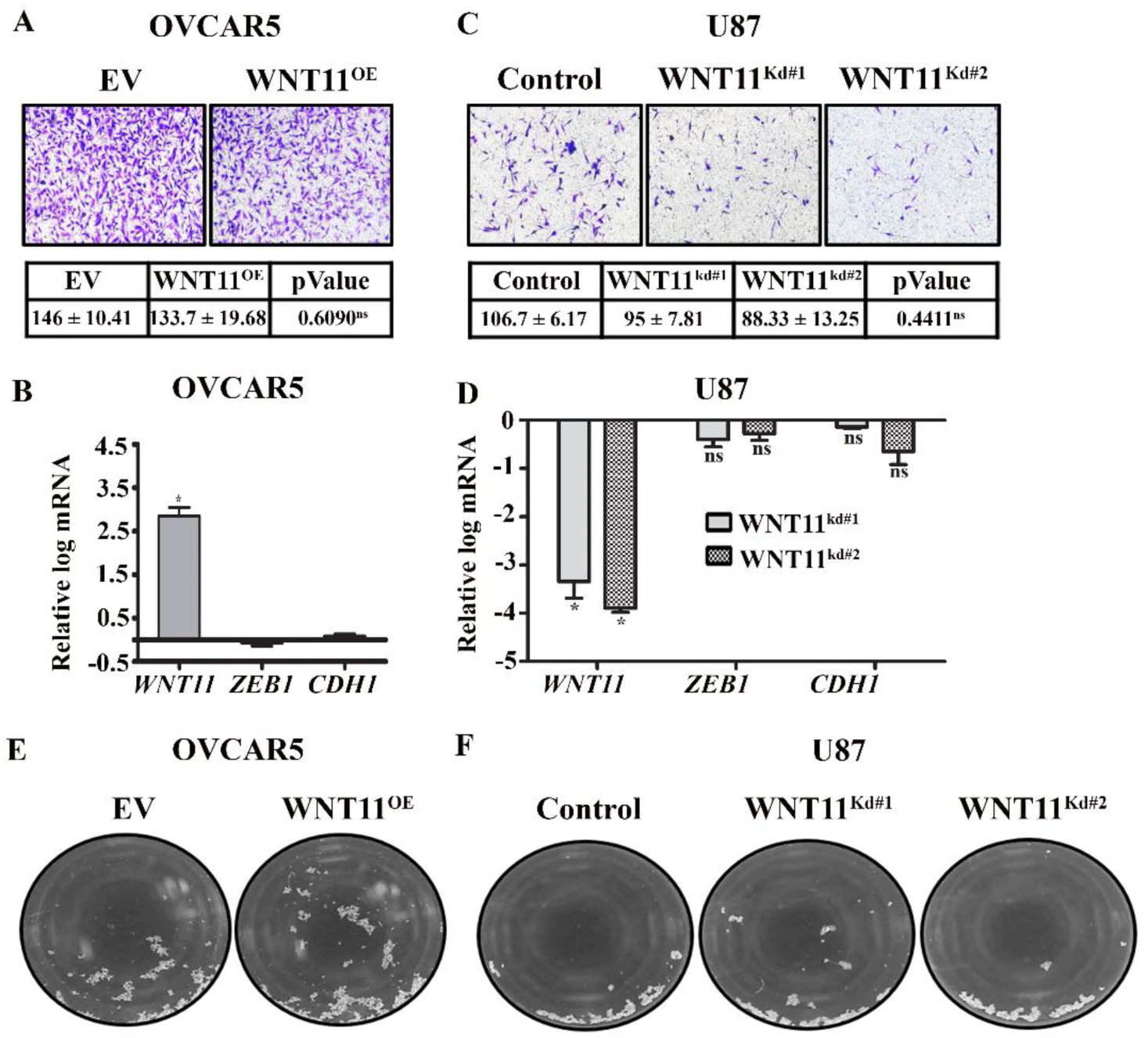
Study the impact of WNT11 on the invasiveness and sphere growth of OVCAR5 and U87 cells. **(A)** Representative images of matrigel invasion assays performed on OVCAR5 with or without WNT11 overexpression. Quantification of invading cells is shown below the images and presented as mean ± SEM, with corresponding p values indicating statistical significance. **(B)** Quantitative PCR analysis of key epithelial-mesenchymal transition (EMT) genes in the same cells as **(A)**; baseline expression is defined by the EV control cells which is set at 0. **(C)** Representative images of matrigel invasion assays performed on U87 cells with or without WNT11 knockdown. Quantification of invading cells is shown below the images and presented as mean ± SEM, with corresponding p values indicating statistical significance. **(D)** Quantitative PCR analysis of EMT gene expression in response to WNT11-knockdown in U87 cells. Data are presented as mean ± SEM (n ≥ 3, n is the number of biological replicates). *p < 0.05 or ns (not significant) are compared to control cells which set the baseline. **(E, F)** Representative microscopic images of third generation spheres from serial replating assays on OVCAR5 cells with or without WNT11 overexpression **(E)** and on U87 cells with or without WNT11 knockdown **(F)**. EV, Empty Vector control.

In our recent study that identified WNT11 as a response gene to RAB4A, we also identified RAC1 as a critical mediator of RAB4A regulation of EMT and stemness. We therefore evaluated RAC1 activity in response to WNT11 manipulation in both WNT11 responsive and un-responsive cell lines. Interestingly, in the WNT11-low responsive MDA-MB-231, PC3 and SNB19 cells, exogenous expression of WNT11 led to suppression of RAC1 activity that was high at baseline (Fig. 5A-C). Conversely, WNT11 knockdown in WNT11-high responsive MCF7 and H1299 cells resulted in a marked increase in RAC1 activation from low baseline (Fig. 5D-E). These results from WNT11 responsive cells clearly showed that WNT11 is a negative regulator of RAC1 activation. We performed similar studies on the WNT11 unresponsive OVCAR5 and U87 cells. Strikingly, manipulating WNT11, either by over-expression in OVCAR5 or knockdown in U87 cells, did not alter the RAC1 activity (Fig. 5F-G); in WNT11-low OVCAR5 cells, the RAC1 activity remained high and in WNT11-high U87 cells the RAC1 activity was persistently undetectable. These findings suggest that for the WNT11-unresponsive cells, RAC1 activation is uncoupled from WNT11 expression. Together, these data suggest that WNT11 modulates RAC1 activity in a context-dependent manner and raises the interesting possibility that the functional impacts of WNT11 modulation on cancer cell invasion and stemness may depend on the uninterrupted chain of signaling from WNT11 to RAC1.

**Figure 5.**
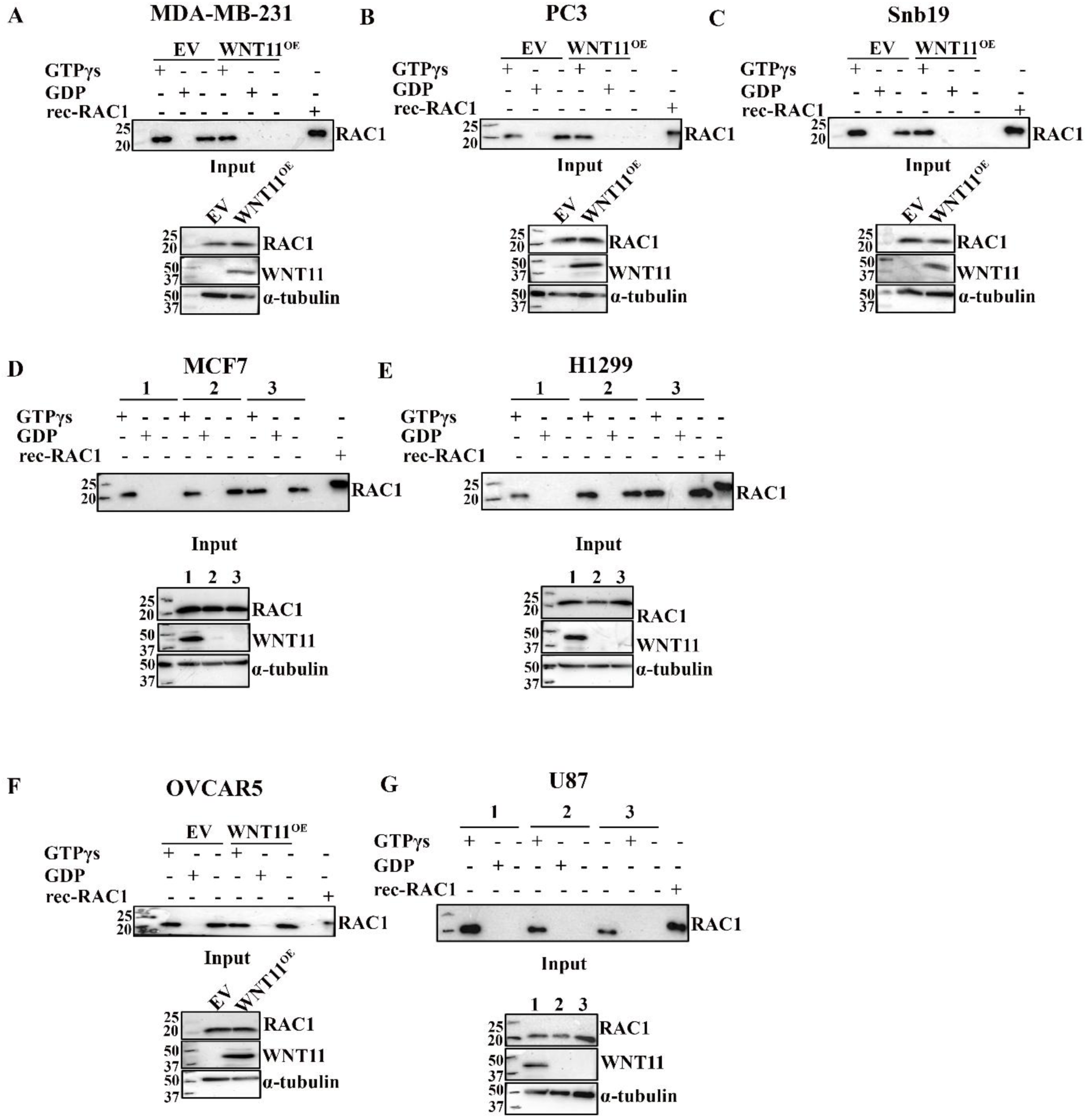
Influence of WNT11 on cancer stemness and invasion and its ability to modulate RAC1 activity. **(A–C)** Pulldown assays assessing GTP-bound (active) RAC1 in response to WNT11 overexpression in the WNT11 responsive WNT11-low cancer cell lines. Each panel includes: Lane 1 – lysate incubated with GTPγS (positive control); Lane 2 – lysate incubated with GDP (negative control); Lane 3 – untreated lysate (endogenous RAC1 activity). Lower panels show input blots for WNT11 and RAC1. **(D–E)** RAC1 activation in response to WNT11 knockdown in WNT11 responsive WNT11-high cell lines. The numeric label indicates the lysates were prepared from cells expressing control shRNA (1); Wnt11 shRNA#1 (2) and WNT11 shRNA#2 (3). At the bottom of the panel is a blot of the lysate without a pull-down. α-Tubulin was the loading control. **(F-G)** RAC1 activation in response to WNT11 modulation in WNT11 unresponsive cell lines. EV, Empty Vector control.

### Direct modulation of RAC1 activity alters invasion, EMT and stemness in WNT11-un-responsive cells

Given the observed association between WNT11 regulation of RAC1 activity and phenotypical impact, we hypothesized that direct manipulation of RAC1 activation in WNT11 unresponsive cells would lead to expected changes of cancer phenotypes such as invasion, EMT and stemness. To this end, we introduced a dominant negative RAC1 mutant (RAC1^DN^; T17N) into OVCAR5 cells that have constitutively high RAC1 activity that is not responsive to WNT11 over-expression. The RAC1^DN^ expression significantly reduced invasive capacity (Fig. 6A), reduced third generation sphere formation (Fig. 6B–C), and decreased OCT4, SOX2 and NANOG expression (Fig. 6D), suggesting that direct inactivation of RAC1 has the same impact as WNT11 in responsive cells. In parallel, a constitutively-active RAC1 mutant (RAC1^CA^; Q61L) was introduced into U87 cells, which exhibits persistently low RAC1 activity despite efficient WNT11 knockdown. Consistent with our hypothesis, expression of RAC1^CA^ enhanced invasion (Fig. 6E), third-generation sphere formation (Fig. 6F-G), and upregulated OCT4, SOX2, and NANOG gene expression (Fig. 6H). Taken together, these results demonstrate that direct manipulation of RAC1 activity result in the phenotype changes in the WNT11 unresponsive cells that mimic the effects of WNT11 in responsive cells, supporting the conclusion that RAC1 is a necessary and sufficient mediator of WNT11 in its role as a tumor suppressor for epithelial cancers.

**Figure 6.**
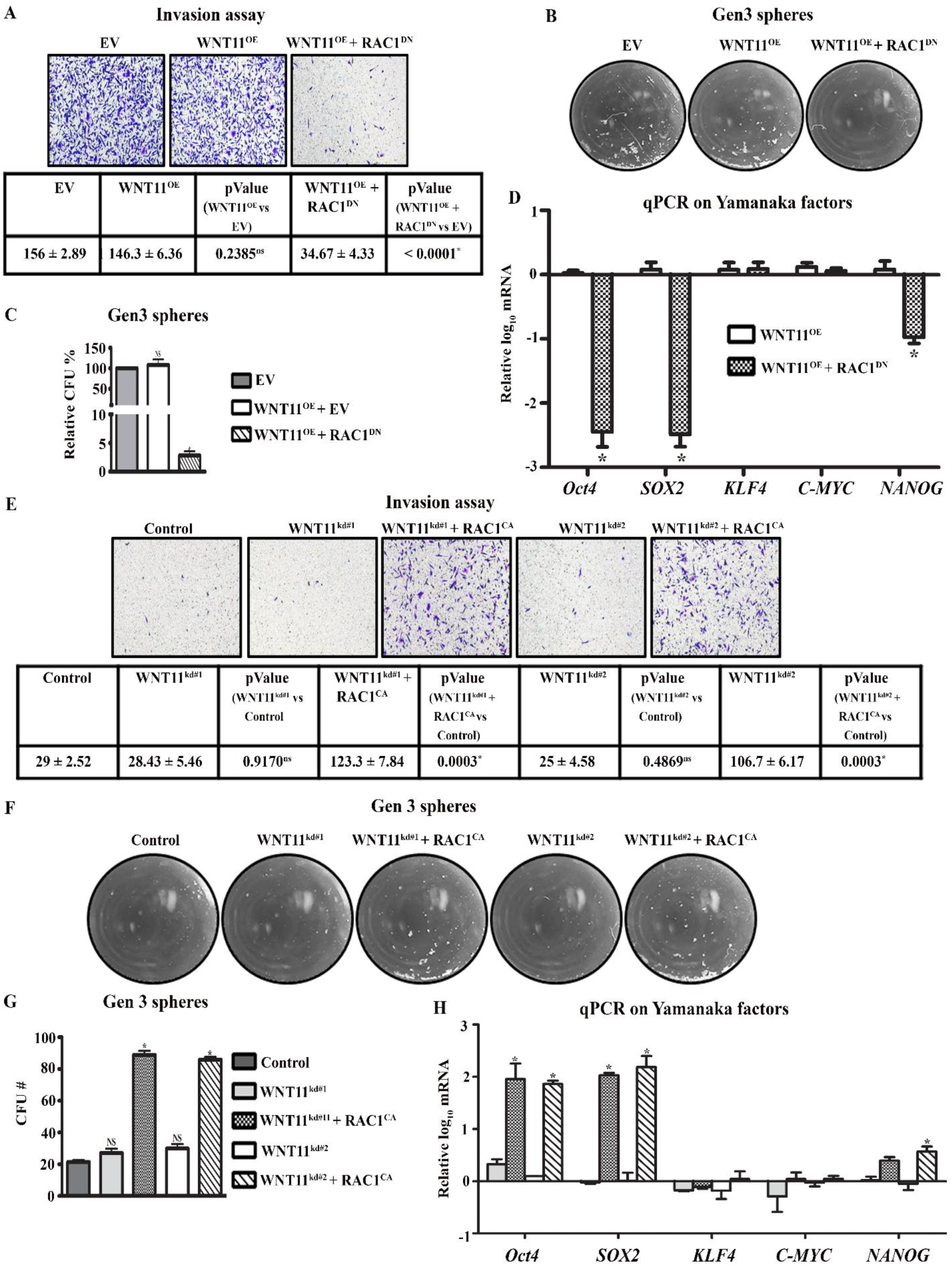
The impact of direct modulation of RAC1 activity on invasion, EMT and stemness in WNT11-unresponsive cells. **(A)** Representative images from matrigel invasion assays on OVCAR5 cells stably overexpressing WNT11, with or without the expression of a dominant-negative RAC1 mutant (RAC1^DN^). Quantification of invading cells is shown below the images and presented as mean ± SEM, with corresponding p values indicating statistical significance. **(B)** Representative images of third generation spheres in serial replating sphere formation assays using the same cell lines as in **(A)**. **(C)** Quantification of third-generation (Gen-3) spheres from the experiment shown in **(B)**. **(D)** qPCR analysis of Yamanaka stemness and NANOG gene expression using the cells in **(A–C)**. Data are presented as mean ± SEM (n ≥ 3, n is number of biological replicates). *p < 0.05 compared to control cells, set as baseline. **(E)** Representative images from matrigel invasion assays on U87 cells with stable WNT11 knockdown, with or without co-expression of constitutively active RAC1 (RAC1^CA^). Quantification of invading cells is shown below the images and presented as mean ± SEM, with corresponding p values indicating statistical significance. **(F)** Representative images of third generation spheres from serial replating sphere formation assays using the same cell line as **(E)**. **(G)** Quantification of third-generation (Gen-3) spheres from the experiment shown in **(F)**. **(H)** qPCR analysis of Yamanaka stemness gene and NANOG expression in the cells as in **(E-G)**. Data are presented as mean ± SEM (n ≥ 3, n is the number of biological replicates). *p < 0.05 compared to control cells, set baseline. EV, Empty Vector control.

## DISCUSSION

Non-canonical WNT signaling has emerged as a critical regulator of tumor cell plasticity, yet it remains far less explored than its canonical counterpart [4]. Among ligands involved in non-canonical activity, WNT11 has drawn interest due to its dual and often contradictory roles across different cancers. Although WNT11 has been linked to enhanced invasiveness in colorectal and pancreatic cancers [28,29], other evidence suggests that it suppresses β-catenin signaling and promotes differentiation, implying tumor-suppressive functions [30,31]. These divergent roles underscore the need for mechanistic dissection of context-dependent actions of WNT11 in human cancers.

Several studies have reported pro-tumorigenic roles of WNT11, particularly on its regulation of cancer cell migration, invasion, and metastatic behavior. In prostate cancer, WNT11 has been shown to promote cell motility, invasion, and neuroendocrine differentiation, phenotypes associated with aggressive disease progression [32,33]. Similarly, in pancreatic ductal adenocarcinoma, elevated WNT11 expression was shown to correlates with enhanced invasiveness and poorer clinical outcome [28]. Several breast cancer studies report the pro-invasive and pro-metastasis role of WNT11 [34]. One study report that WNT11 binding to ROR2 transmits signaling to RHO/ROCK pathways to promote tumor cell invasion [35]. Another study reports WNT11-dependent activation of Wnt–planar cell polarity (PCP) signaling enhances breast cancer cell metastatic behavior through exosome-mediated tumor microenvironment regulation [34]. In colorectal cancer, WNT11 has been implicated in invasion downstream of tumor-secreted factors such as AGR2 and non-canonical CaMKII/JNK signaling pathways [36]. In contrast to these studies, our findings demonstrate that WNT11 suppresses stemness, invasion and tumor initiation in a panel of epithelial cancer models in which WNT11 signaling remains functionally coupled to RAC1 inhibition.

It is important to point out that, rather than contradicting the prior reports on WNT11 acting as an oncogene, this study underscores the complex and context-dependent signaling and function of WNT11 in cancer. Mechanistically, the study brings into focus the notion that the ability of WNT11 to act as a tumor suppressor is dependent on whether it can negatively regulate RAC1 activity. In the complex network of WNT signaling, the availability of every component of signaling chain and crosstalk pathways - from cell surface receptor, scaffold/adaptors to downstream effectors such as RHO family GTPases and their regulators can contribute to the biological outcome. Hence, the specific cell context of each cancer cell type dictates the ultimate role of WNT11, either as tumor suppressor or oncogene. In the study, we have evaluated multiple cancer cell lines across tissue types and clarified that WNT11 acts as tumor suppressor through RAC1 inactivation. A better understanding of the specific context and signaling circuitry is critical for potentially targeting WNT11 in selected group of cancers. Hence, further investigation is necessary to delineate the molecular framework to predict WNT11-mediated cancer phenotypes.

In addition to the ability of WNT11 to engage RAC1 that can account for the context-dependence of responsive and non-responsive cancers, there are additional interesting observations in this regard. One of these is that, while WNT11 manipulation led to robust changes in the expression of five key stem regulators we surveyed – OCT4, SOX2, KLF4, C-MYC and NANOG – manipulating RAC1 activation only engaged three of these five: OCT4, SOX2 and NANOG. Hence, it is reasonable to speculate that there are likely other effectors downstream of WNT11 that regulate KLF4 and C-MYC.

We observed that parental MCF7 cells failed to form palpable tumors in NOD-SCID mice within the observation period without estrogen stimulation. This is consistent with the established understanding that the ER+ breast cancer MCF7 cells are estrogen receptor-positive and exhibit hormone-dependent growth in xenograft models [37–39]. It is interesting that WNT11 depletion abolished this estrogen-dependency and promoted tumor formation. In the case of non-small cell lung cancer cell line H1299, these cells have diminished tumor forming ability unless going through the cell selection for CD44+ or stem cell enrichment protocol first before implanting into immune compromised mice [24–27,40]. However, WNT11 knockdown conferred the parental H1299 cells with tumor forming capacity. These findings demonstrate that WNT11 can function as a potent suppressor of tumor initiation across multiple epithelial cancer models when coupled to RAC1 regulation.

As WNT signaling plays major roles in cancer stem cells, the development of inhibitors of its signaling pathways has garnered much attention in the past two decades [17–20,41,42]. Development of comprehensive stratification strategies and new ligand/receptor-specific drugs demands better understanding of individual WNT ligand signaling in different cellular contexts, as exemplified here by WNT11. The findings in this study provide new insights into the non-canonical WNT signaling landscape and position WNT11 and its downstream effectors - particularly RAC1 - as functionally significant modulators of cancer cell plasticity and thereby promising targets for therapeutic intervention in aggressive cancers.

## MATERIALS & METHODS

### Cancer cell lines and culture

Human breast cancer cell lines (MDA-MB-231, MCF7), prostate cancer cells (PC3), non-small cell lung cancer cells (H1299), glioblastoma cells (SNB19, U87), and ovarian cancer cells (OVCAR5) were obtained from ATCC (Rockville, MD) and confirmed to be mycoplasma-negative. Cells were cultured in DMEM (Nacalai, USA) supplemented with 10% Fetal Bovine Serum (FBS) and penicillin/streptomycin (Hyclone, IL, USA) under standard conditions. Actinomycin D (Thermo Fisher, USA) treatments were carried out using established protocols. All cell lines used in this study, both parental and the derived stable lines (parental, EV, non-target shRNA control, WNT11 overexpressing and WNT11 knockdown) were authenticated by 1st BASE (Axil Scientific, Singapore) using the GenePrint® 24 System (Promega), which amplifies 24 STR loci and the Amelogenin gender-determining locus [43]. All STR profiles (submitted as supplementary data) matched reference databases (ATCC/DSMZ).

### Generation of stable cell lines and transient transfection

Lentiviral particles were generated in HEK293T cells using Lipofectamine 2000 (Invitrogen, USA) per standard lab protocols [44]. Stable WNT11 knockdown was achieved using shRNAs cloned into pLL3.7 vectors (Supplementary Table S1A). Cells were transduced with lentiviral particles encoding specific shRNAs and the control (non-target shRNA) in the presence of 8 μg/mL polybrene (Sigma-Aldrich). After 24 h, cells were selected using puromycin (2 µg/mL) for 5–7 days. Knockdown efficiency was validated by qPCR and Western blotting in comparison to the stable control (non-target shRNA) (Supplementary Table S1A). Full length WNT11 was amplified from MCF7 cDNA using primers and the coding sequence (Supplementary table S1B) was cloned into pMSCV vector for overexpression. Retroviral particles were produced as described above and the transduced cells were selected using blasticidin (20 µg/mL) for 15 – 20 days. Constructs expressing dominant-negative (T17N) or constitutively active (Q61L) RAC1 mutants were prepared as described previously [16] and used in selected cell lines. For live-cell imaging, cells were seeded in 96-well plates and imaged using the IncuCyte ZOOM system (Essen Bioscience, USA) at indicated time points [45].

### Quantitative PCR (qPCR)

RNA was isolated using the Tissue Total RNA Mini Kit (Favorgen Biotech), and cDNA was synthesized with ReverTra Ace qPCR RT Master Mix (Toyobo). qPCR was performed using SYBR Green (Roche) on a CFX96 instrument (Bio-Rad). Primers are listed in Supplementary Table S1C. Cancer stemness gene profiling was performed according to the manufacturer’s protocol using the SAB-targeted multiplex array (Bio-Rad, USA). Heatmaps were generated using Morpheus software.

### Immunoblotting and RAC1 pull-down assay

Whole-cell lysates were prepared in RIPA buffer (50 mM Tris, pH 7.6, 150 mM NaCl, 1% Triton X-100, 0.1% SDS) containing protease and phosphatase inhibitors (#4693159001, Roche, WI, #P0044, Sigma-Aldrich, St. Louis, MO), and processed using lab standard protocol [46]. Antibodies used for the blotting are listed in Supplementary Table S1D. RAC1 activation assays were performed using a pull-down assay kit to detect GTP-bound RAC1, following the manufacturer’s protocol from Cytoskeleton, Inc. (BK035; Denver, CO). Total RAC1 and active RAC1 levels were detected by immunoblotting as described above.

For analysis of secreted WNT11, media were collected from relevant MDA-MB-231 derived cells. Cells were seeded at 2 × 10⁶ per 10-cm plate and incubated for 24 h. After incubation, medium was replaced with serum-free DMEM, and conditioned media were collected after another 24 h followed by centrifugation at 1500 rpm for 5 min to get the clear media, which were concentrated using 10 kDa cut-off ultrafiltration columns (Amicon, Merck Millipore, USA) at 4000 g for 45 min at 4°C until the volume was reduced to ∼150 μl. For deglycosylation, the concentrated samples were treated with NEB Protein Deglycosylation Mix II per manufacturer’s instructions (NEB, USA). WNT11 was visualized by SDS-PAGE and immunoblot. Parallel whole-cell lysates were collected and used for comparison.

### Sphere formation assay

Sphere formation efficiency was analyzed by seeding 400 cells/well in ultra-low attachment 96-well plates (Corning) using DMEM-F12 supplemented with 0.5% methylcellulose, B-27, and N2 (Gibco). Spheres were cultured for 10–14 days [47]. For serial passaging, spheres were dissociated using Accutase (Gibco) and re-seeded under identical conditions [47]. Sphere numbers were quantified using OpenCFU software.

### Invasion assays

Transwell invasion assays were conducted using 24-well inserts with 8.0 μm pores (#3422, Corning, USA). Inserts were coated with a diluted Matrigel solution and allowed to solidify overnight. Following serum starvation, 1 × 10⁵ cells suspended in 100 μl serum-free medium with 0.1% BSA were added to the upper chamber. The lower chamber contained 0.5 ml of medium with 10% FBS. After 24 h incubation at 37°C, membranes were fixed with 4% paraformaldehyde, stained with 0.2% crystal violet (in MeOH), and non-invaded cells were removed [48]. Invasion was quantified microscopically. For invasion assays using WNT11 conditioned media, the conditioned media from WNT11 overexpressing MDA-MB-231 cells was collected as described under “Immunoblotting methods” and applied to the upper chamber containing control cells. After 24 h, cells were fixed, stained, and invasion quantified to assess paracrine activity of secreted WNT11. Quantification of invading cells was performed by counting stained cells using ImageJ.

### Animals and xenografts

All animal protocols were approved by the SingHealth Institutional Animal Care and Use Committee (IACUC protocol no. 2021/SHS/1627). Eight-week old female NOD-SCID mice were purchased from InVivos (InVivos Pte Ltd, Singapore) and housed under pathogen-free conditions with a 12 h light/12 h dark cycle and provided food and water ad libitum. For subcutaneous xenografts, cells were harvested using trypsin, counted, and assessed for viability by trypan blue exclusion counting. Only cell preparations with >95% viable cells were used for implantation. A total of 5 × 10⁶ cells were suspended in 100 µL of a 1:1 mixture of PBS and Matrigel (BD Biosciences) and injected subcutaneously into the flank of each mouse; no anesthesia was required for this procedure. At least five mice (N = 5) were used for each condition. Tumor growth was monitored twice weekly using calipers. Mice were euthanized via CO_2_ inhalation followed by cervical dislocation when tumors reached 1 cm³ or at the experimental endpoint (90 days), in accordance with guidelines and regulations of SingHealth IACUC.

### Statistical analysis

All data represent mean ± SEM from at least three independent experiments. Statistical analyses were performed using GraphPad Prism. Significance was evaluated using unpaired t-tests, one-way or two-way ANOVA with appropriate post hoc tests (Tukey’s or Dunnett’s). A p-value < 0.05 was considered statistically significant.

## Supporting information

Cell Identity Confirmation

## Acknowledgements

We appreciate the support from Duke-NUS core facility and vivarium.

## Funding

The work is supported by National Medical Research Council Individual Research Grant (MOH-000944-01) and the Duke-NUS Medical School Block Fund.

## Author contribution statement

S.K designed and performed research, analyzed data and wrote the manuscript; P.J.C provided expertise and edited the manuscript; M.W. conceived and supervised the study, interpreted the data, wrote and revised the manuscript, and provided funding support and ethical protocols. All authors reviewed and approved the manuscript.

## Competing Interests

The authors declare no competing interests.

## Data Availability Statement

The data supporting the findings of this study are available within the article and its supplementary information files. Additional data are available from the corresponding author upon reasonable request.

## Notes

### Competing Interest Statement

The authors have declared no competing interest.

