## Supplementary Figures for "WNT11 suppresses tumor initiation and invasion by inactivating RAC1"

### Figure S1.

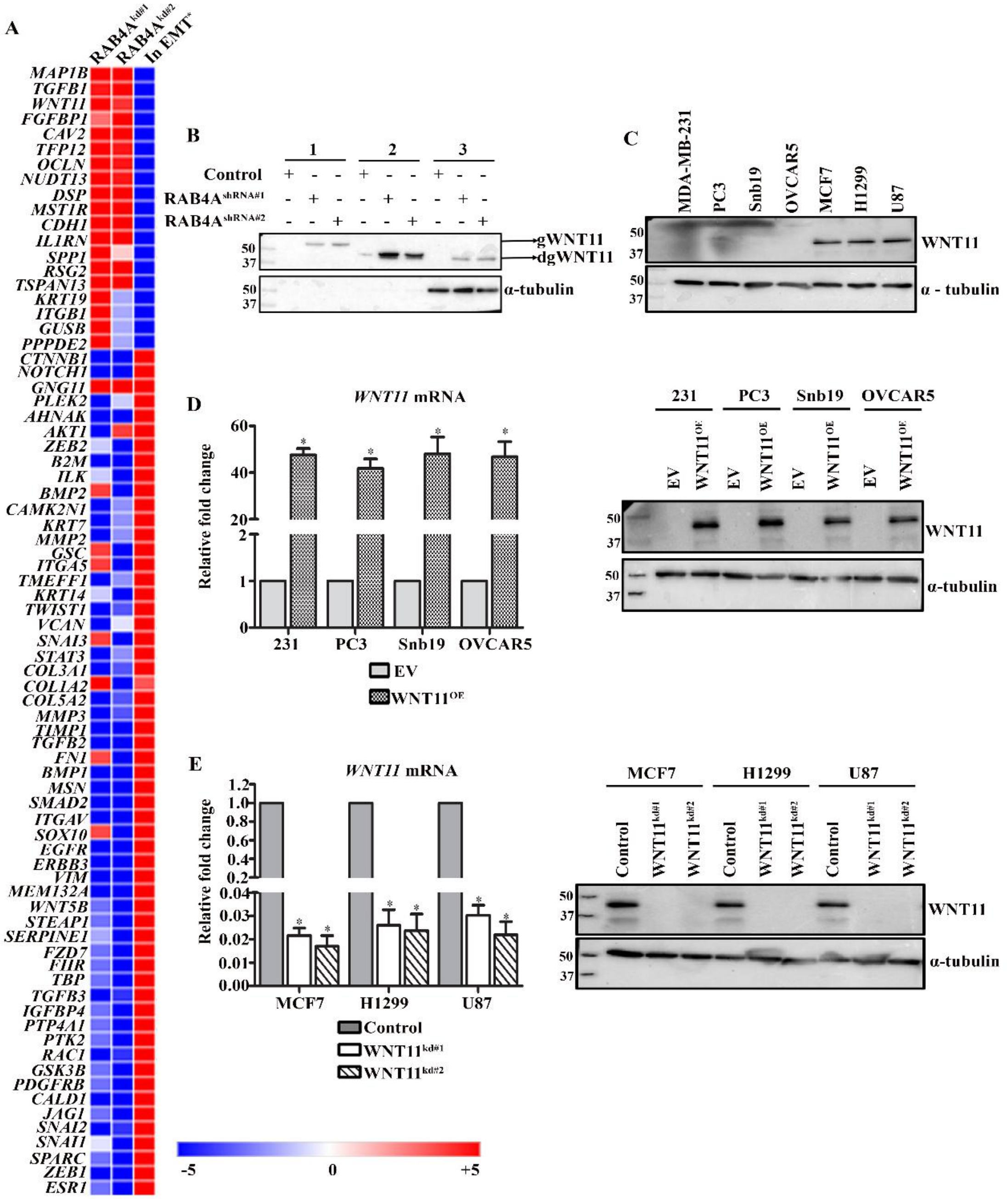


**Figure S1. WNT11 expression in cancer.**
**(A)** Multiplex qPCR analysis of EMT gene expression changes in response to RAB4A knockdown in MDA-MB-231 cells as previously published [1]. The heat map of the first two column display Z-Scores derived from normalized qPCR data in RAB4A knockdown clones (Kd#1 and Kd#2) relative to control cells; the third column shows the expected change of the individual gene expression level when the cell undergoes canonical EMT process. **(B)** Immunoblot analysis of WNT11 level in media collected from MDA-MB-231 cells with stable RAB4A knockdown. Group

1: concentrated media; group 2: concentrated media underwent deglycosylation treatment; group 3: cell lysate. **(C)** Immunoblot analysis screening for WNT11 expression in a panel of human cancer cell lines; α-Tubulin serves as the loading control. **(D, E)** Validation of WNT11 modulation in human cancer cell lines. Quantitative PCR analyses and immunoblot confirming stable WNT11 overexpression (D – left & right) and knockdown (E – left & right), demonstrating robust changes in WNT11 expression protein and mRNA levels. α-Tubulin serves as the loading control. EV represents "Empty Vector" as expression control.

Figure S2

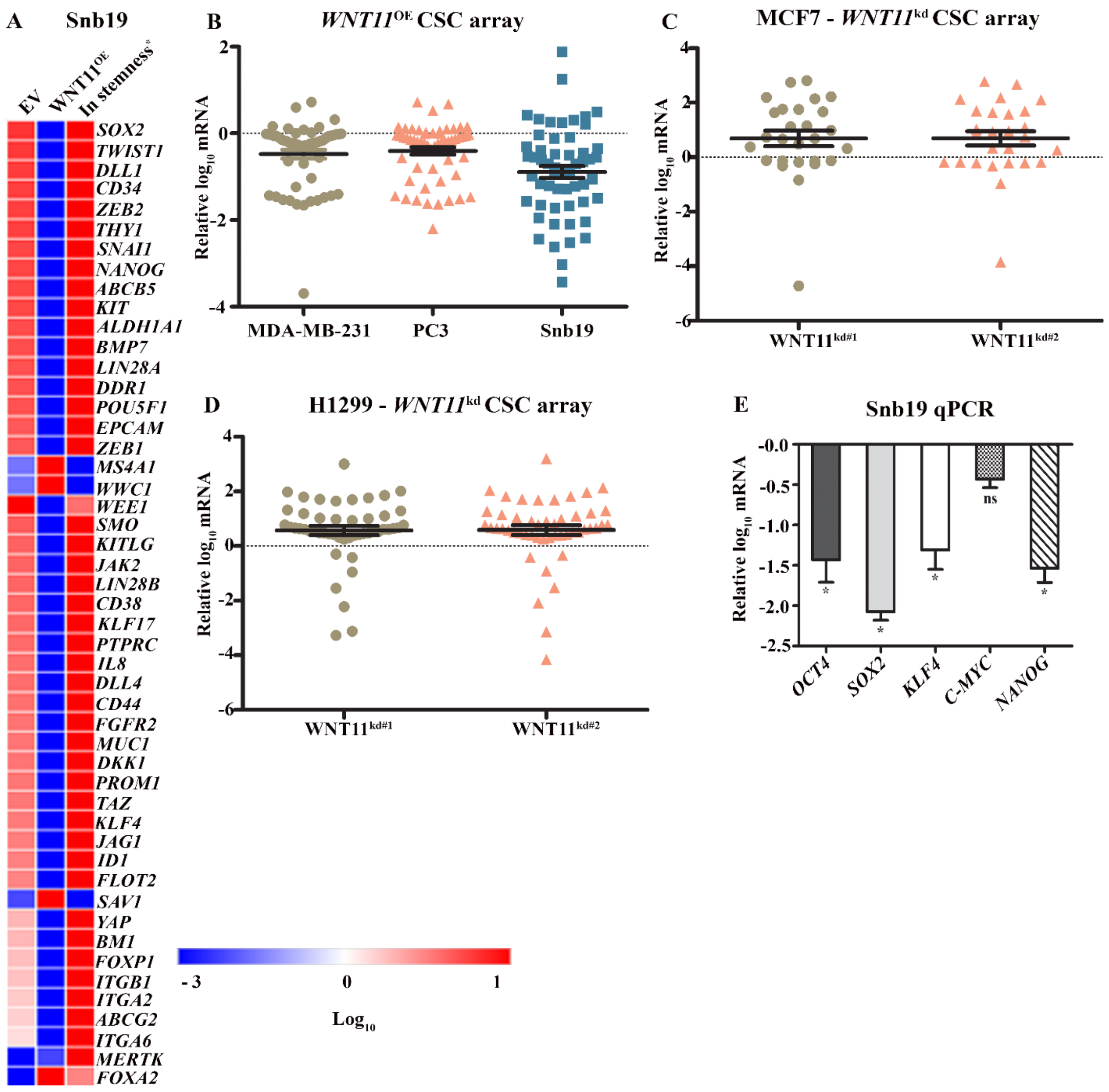


**Figure S2. Stemness gene expression analysis in response to WNT11 modification.** Multiplex qPCR was performed on 87 genes implicated in cancer stemness. Relative expression was calculated against control cells. **(A)** Genes that were changed in response to WNT11 expression in Snb19 cells. Grouped scatter plots are shown for genes altered by ≥1-fold compared to respective controls. **(B)** Data from cells with WNT11 overexpression. **(C, D)** Data from cells stably expressing WNT11 shRNA. **(E)** qPCR analysis for the expression of Yamanaka Factors and NANOG in same cells as in **(A)**. Data are presented as mean ± SEM ($n \geq 3$; n is the number of biological repeats). $*p < 0.05$ compared to control cells, which are set as the baseline.”

**Figure S3.**

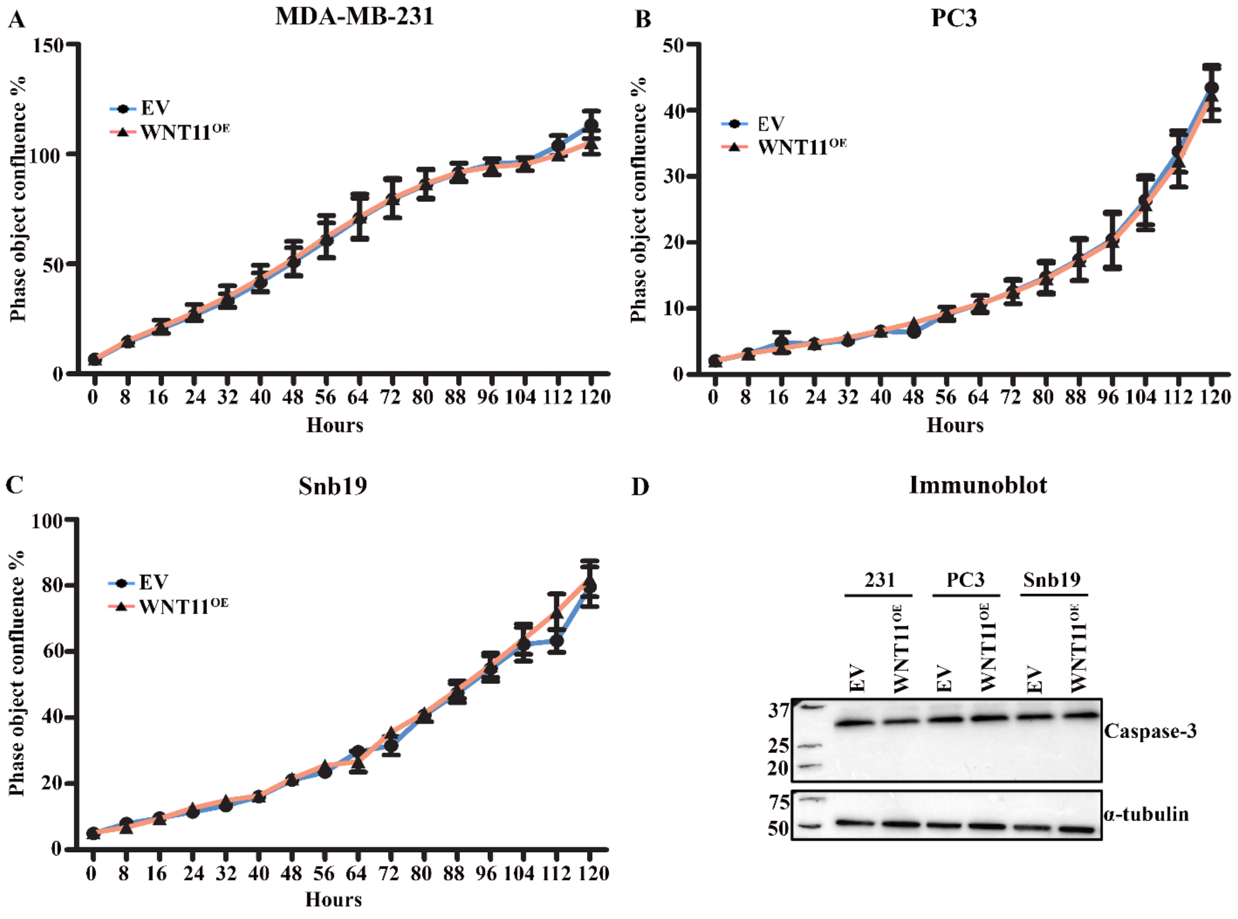


**Figure S3. Exogenous expression of WNT11 in WNT11-low cancer cells does not affect their proliferation or viability under the adherent culturing condition.**

**(A-C)** Proliferation of MDA-MB-231 **(A)**, PC3 **(B)**, and SNB19 **(C)** cells stably overexpressing WNT11 under adherent culture conditions, compared to empty vector control cells. Proliferation was monitored continuously for 120 hours using IncuCyte live-cell imaging.

**(D)** Immunoblot analysis of caspase-3 expression to assess cell viability. α-Tubulin serves as the loading control.

EV represents "Empty Vector" as expression control.

**Figure S4**

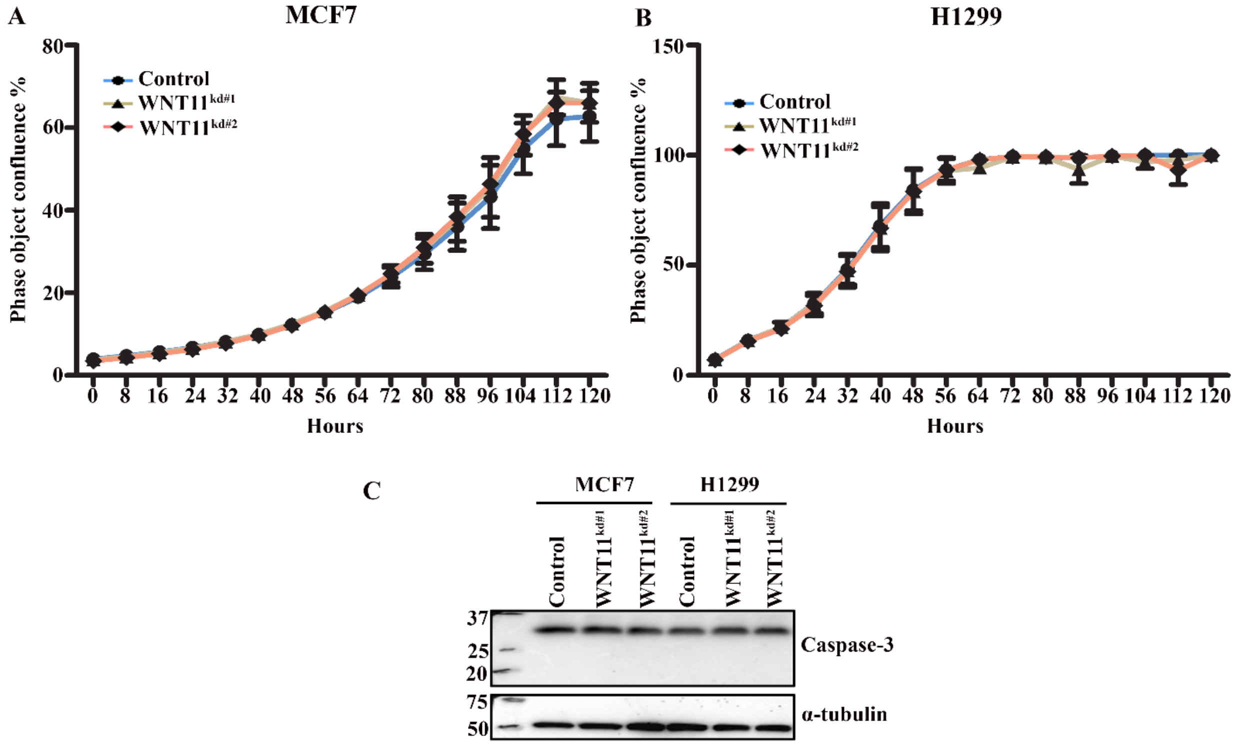


**Figure S4. Stable knockdown of WNT11 in WNT11-high cancer cells does not change their proliferation or viability under the adherent culturing condition.**

**(A-B)** Proliferation of stable WNT11 knockdown MCF7 and H1299 cells under adherent culture conditions, compared to cells expressing control shRNA. Proliferation was monitored continuously for 120 hours using IncuCyte live-cell imaging.

**(C)** Immunoblot analysis of caspase-3 expression to assess cell viability. α-Tubulin serves as the loading control.

### Supplementary Tables

**Supplementary Table S1A –** WNT11 shRNA sequenced cloned into pLL3.7

| S.No | Description | Sequence (5'-3') |
|---|---|---|
| 1 | CDS | GCCAATAAACTGATGCGTC |
| 2 | UTR | GCTTGTGCTTTGCCTTCAC |
| 3 | Control (Non-target shRNA) | GCGCGATAGCGCTAATAAT |

**Supplementary Table S1B** – Primer sequence used to clone WNT11 cDNA into pMSCV and the cloned WNT11 CDS sequence

| S.No | Forward primer (5'-3') | Reverse primer (5'-3') |
|---|---|---|
| 1 | ATGAGGGCGCGGCCGCA | TCACTTGCAGACATAGCGCTCCA |
| **Cloned WNT11 CDS sequence** | | |
| ATGAGGGCGCGGCCGCAGGTCTGCGAGGCGCTGCTCTTCGCCCTGGCGCTCCAGA<br>CCGGCGTGTGCTATGGCATCAAGTGGCTGGCGCTGTCCAAGACACCATCGGCCCT<br>GGCACTGAACCAGACGCAACACTGCAAGCAGCTGGAGGGTCTGGTGTCTGCACAG<br>GTGCAGCTGTGCCGCAGCAACCTGGAGCTCATGCACACGGTGGTGCACGCCGCCC<br>GCGAGGTCATGAAGGCCTGTCGCCGGGCCTTTGCCGACATGCGCTGGAACTGCTC<br>CTCCATTGAGCTCGCCCCCAACTATTTGCTTGACCTGGAGAGAGGGACCCGGGAG<br>TCGGCCTTCGTGTATGCGCTGTCGGCCGCCGCCATCAGCCACGCCATCGCCCGGGC<br>CTGCACCTCCGGCGACCTGCCCGGCTGCTCCTGCGGCCCCGTCCCAGGTGAGCCA<br>CCCGGGCCCGGGAACCGCTGGGGAGGATGTGCGGACAACCTCAGCTACGGGCTCC<br>TCATGGGGGCCAAGTTTTCCGATGCTCCTATGAAGGTGAAAAAAACAGGATCCCA<br>AGCCAATAAACTGATGCGTCTACACAACAGTGAAGTGGGGAGACAGGCTCTGCGC<br>GCCTCTCTGGAAATGAAGTGTAAGTGCCATGGGGTGTCTGGCTCCTGCTCCATCCG<br>CACCTGCTGGAAGGGGCTGCAGGAGCTGCAGGATGTGGCTGCTGACCTCAAGACC<br>CGATACCTGTCGGCCACCAAGGTAGTGCACCGACCCATGGGCACCCGCAAGCACC<br>TGGTGCCCAAGGACCTGGATATCCGGCCTGTGAAGGACTCGGAACTCGTCTATCT<br>GCAGAGCTCACCTGACTTCTGCATGAAGAATGAGAAGGTGGGCTCCCACGGGACA<br>CAAGACAGGCAGTGCAACAAGACATCCAACGGAAGCGACAGCTGCGACCTTATG<br>TGCTGCGGGCGTGGCTACAACCCCTACACAGACCGCGTGGTCGAGCGGTGCCACT<br>GTAAGTACCACTGGTGCTGCTACGTCACCTGCCGCAGGTGTGAGCGTACCGTGGA<br>GCGCTATGTCTGCAAGTGA | | |

**Supplementary Table S1C** – qPCR primers

| Gene | Forward primer (5'-3') | Reverse primer (5'-3') |
|---|---|---|
| WNT11 | AGGACTCGGAACTCGTCTAT | TGTTGCACTGCCTGTCTT |
| SOX2 | GCTACAGCATGATGCAGGACCA | TCTGCGAGCTGGTCATGGAGTT |
| 4-Oct | CAGGAGATATGCAAAGCAGAAACCCT | TCGGGCACTGCAGGAACAAATTC |
| KLF4 | GCGAACCCACACAGGTGAGAAAC | ACGGTAGTGCCTGGTCAGTTCATC |
| C-MYC | GCAGCGACTCTGAGGAGGAACA | TGGCCTCCAGCAGAAGGTGATC |

| | | |
|---|---|---|
| NANOG | GAACCTCAGCTACAAACAGGTGAAGA | TCCCTGCGTCACACCATTGCTATTC |
| ZEB1 | AGTGGTCATGAAAATGGAAC | AGGTGTAACTGCACAGGGAGC |
| CDH1 | GTCCTGGGCAGAGTGAATTT | GACCAAGAAATGGATGTGTGG |

**Supplementary Table S1D** – Antibodies

| Antibody | Cat No | Manufacturer |
|---|---|---|
| WNT11 | GTX105971 | Genetex |
| Caspase-3 | 9662 | Cell Signalling |
| α - tubulin | T9026 | Sigma Aldrich |
