## Supplementary material for "WNT11 suppresses tumor initiation and invasion by inactivating RAC1": Cell Identity Confirmation

Axil Scientific Pte Ltd
2 Tukang Innovation Grove, #06-01, JTC MedTech Hub,
Singapore 618305
T: +65 6775 7318
F: +65 6775 7211
E:

Apical Scientific Sdn Bhd
Lot 7-1 to 7-4, Jalan SP 2/7, Taman Serdang Perdana,
Seksyen 2, 43300 Seri Kembangan, Selangor, Malaysia
T: +603 8943 3252
F: +603 8943 3243
E:

1st BASE
YOUR RESEARCH, OUR BUSINESS
www.base-asia.com

### CUSTOMER INFORMATION

| | | | |
|---|---|---|---|
| Name | Subbulakshmi Karthikeyan | Order ID | 1548 |
| Address | | Date Sample Received | 2-Feb-2026 |
| Email | | Report Date | 5-Feb-2026 |

### METHODOLOGY

Twenty-four short tandem repeat (STR) loci plus the gender determining locus, Amelogenin, were amplified using the commercially available GenePrint® 24 System from Promega. The sample was processed using the Applied Biosystems™ DNA Analyzer. Data were analyzed using GeneMapper® v4.0 software (Applied Biosystems™). Appropriate positive and negative controls were run and confirmed for each sample submitted.

### SAMPLE INFORMATION

| **Sample Name** | **Cell Line Designation** |
|---|---|
| 231 | MDA-MB-231 |

### STR PROFILING RESULTS

| | **Test Result for Sample** | | | | **ExPASy Reference Database Profile** | |
|---|---|---|---|---|---|---|
| **LOCI** | **231** | | | | **MDA-MB-231** | |
| Amelogenin | X | | | | X | |
| D3S1358 | 16 | | | | | |
| D1S1656 | 15 | 17 | | | | |
| D2S441 | 14 | 15 | | | | |
| D10S1248 | 14 | 16 | | | | |
| D13S317 | 13 | | | | 13 | |
| Penta E | 11 | | | | | |
| D16S539 | 12 | | | | 12 | |
| D18S51 | 16 | | | | | |
| D2S1338 | 20 | 21 | | | | |
| CSF1PO | 12 | 13 | | | 12 | 13 |
| Penta D | 11 | 14 | | | | |
| TH01 | 7 | 9.3 | | | 7 | 9.3 |
| vWA | 15 | 18 | | | 15 | 18 |
| D21S11 | 30 | 33.2 | | | | |
| D7S820 | 8 | 9 | | | 8 | 9 |
| D5S818 | 12 | | | | 12 | |
| TPOX | 8 | 9 | | | 8 | 9 |
| DYS391 | | | | | | |
| D8S1179 | 13 | | | | | |
| D12S391 | 17 | 18 | | | | |
| D19S433 | 11 | 14 | | | | |
| FGA | 22 | 23 | | | | |
| D22S1045 | 16 | | | | | |

| | |
|---|---|
| Number of shared alleles between sample and database profile: | 14 |
| Total number of alleles in the database profile: | 14 |
| Percent match between the submitted sample and the database profile: | 100% |

*The allele match algorithm compares the loci highlighted in grey only (8 core loci plus amelogenin).*

### EXPLANATION OF TEST RESULTS

- [ ] The submitted sample profile is human, but not a match for any profile in the STR database.
- [x] The submitted sample profile showed 80% to 100% match for the following ExPASy human cell line(s) in the STR database (8 core loci plus Amelogenin): **MDA-MB-231**
- [ ] The submitted profile is similar to the following ExPASy human cell line(s):
- [ ] The submitted sample is a mixture. Multiple peaks are observed in the STR profiling results.

### ADDITIONAL INFORMATION: Comparative Data Output from ExPASy STR Profile Database

| Accession | Name | Nº Markers | Score | Amel | CSF1PO | D5S818 | D7S820 | D13S317 | D16S539 | TH01 | TPOX | vWA |
|---|---|---|---|---|---|---|---|---|---|---|---|---|
| CVCL_0062 | MDA-MB-231 | 9 | 100.00% | X | 12,13 | 12 | 8,9 | 13 | 12 | 7,9.3 | 8,9 | 15,18 |
| CVCL_YJ26 | MDA-MB-231 shWDR12-4 | 9 | 100.00% | X | 12,13 | 12 | 8,9 | 13 | 12 | 7,9.3 | 8,9 | 15,18 |
| CVCL_VR67 | MDA-MB-231 VIM RFP | 9 | 100.00% | X | 12,13 | 12 | 8,9 | 13 | 12 | 7,9.3 | 8,9 | 15,18 |
| CVCL_JG53 | MDA-MB-231-Luc [JCRB] | 9 | 100.00% | X | 12,13 | 12 | 8,9 | 13 | 12 | 7,9.3 | 8,9 | 15,18 |
| CVCL_VR36 | MDA231-BrM2-831 | 9 | 100.00% | X | 12,13 | 12 | 8 | 13 | 12 | 7,9.3 | 8,9 | 15 |
| CVCL_5998 | MDA231-LM2-4175 | 9 | 100.00% | X | 12,13 | 12 | 8 | 13 | 12 | 7,9.3 | 8,9 | 15 |
| CVCL_VR35 | MDA231-TGL | 9 | 100.00% | X | 12,13 | 12 | 8 | 13 | 12 | 7,9.3 | 8,9 | 15 |
| CVCL_DP48 | MDA-BoM-1833 | 9 | 100.00% | X | 12,13 | 12 | 8 | 13 | 12 | 7,9.3 | 8,9 | 15 |

For alternate database, you may visit https://www.dsmz.de/services/services-human-and-animal-cell-lines/online-str-analysis.html

End of report

**Fragment Analysis**
**- STR Profiling**
**Cell Line Authentication**

Axil Scientific Pte Ltd
2 Tukang Innovation Grove, #06-01, JTC MedTech Hub,
Singapore 618305
T: +65 6775 7318
F: +65 6775 7211
E:

Apical Scientific Sdn Bhd
Lot 7-1 to 7-4, Jalan SP 2/7, Taman Serdang Perdana,
Seksyen 2, 43300 Seri Kembangan, Selangor, Malaysia
T: +603 8943 3252
F: +603 8943 3243
E:

1st BASE
YOUR RESEARCH, OUR BUSINESS
www.base-asia.com

### CUSTOMER INFORMATION

| | | | |
|---|---|---|---|
| Name | Subbulakshmi Karthikeyan | Order ID | 1630 |
| Address | | Date Sample Received | 16-Apr-2026 |
| Email | | Report Date | 23-Apr-2026 |

### METHODOLOGY

Twenty-four short tandem repeat (STR) loci plus the gender determining locus, Amelogenin, were amplified using the commercially available GenePrint® 24 System from Promega. The sample was processed using the Applied Biosystems™ DNA Analyzer. Data were analyzed using GeneMapper® v4.0 software (Applied Biosystems™). Appropriate positive and negative controls were run and confirmed for each sample submitted.

### SAMPLE INFORMATION

| Sample Name | Cell Line Designation |
|---|---|
| 231___EV | MDA-MB-231 |

### STR PROFILING RESULTS

| | Test Result for Sample | | | | ATCC Reference Database Profile | |
|---|---|---|---|---|---|---|
| **LOCI** | **231___EV** | | | | **MDA-MB-231 Breast Adenocarcinoma Human** | |
| Amelogenin | X | | | | X | |
| D3S1358 | 16 | | | | | |
| D1S1656 | 15 | 17 | | | | |
| D2S441 | 14 | 15 | | | | |
| D10S1248 | 14 | 16 | | | | |
| D13S317 | 13 | | | | 13 | |
| Penta E | 11 | | | | | |
| D16S539 | 12 | | | | 12 | |
| D18S51 | 16 | | | | | |
| D2S1338 | 20 | 21 | | | | |
| CSF1PO | 12 | 13 | | | 12 | 13 |
| Penta D | 11 | 14 | | | | |
| TH01 | 7 | 9.3 | | | 7 | 9.3 |
| vWA | 15 | 18 | | | 15 | 18 |
| D21S11 | 30 | 33.2 | | | | |
| D7S820 | 8 | 9 | | | 8 | 9 |
| D5S818 | 12 | | | | 12 | |
| TPOX | 8 | 9 | | | 8 | 9 |
| DYS391 | | | | | | |
| D8S1179 | 13 | | | | | |
| D12S391 | 17 | 18 | | | | |
| D19S433 | 11 | 14 | | | | |
| FGA | 22 | 23 | | | | |
| D22S1045 | 16 | | | | | |

| | |
|---|---|
| Number of shared alleles between sample and database profile: | 14 |
| Total number of alleles in the database profile: | 14 |
| Percent match between the submitted sample and the database profile: | 100% |

*The allele match algorithm compares the loci highlighted in grey only (8 core loci plus amelogenin).*

### EXPLANATION OF TEST RESULTS

☐ The submitted sample profile is human, but not a match for any profile in the STR database.

☑ The submitted sample profile showed 80% to 100% match for the following ATCC human cell line(s) in the STR database (8 core loci plus Amelogenin): **MDA-MB-231 Breast Adenocarcinoma Human**

☐ The submitted profile is similar to the following ATCC human cell line(s):

☐ The submitted sample is a mixture. Multiple peaks are observed in the STR profiling results.

### ADDITIONAL INFORMATION: Comparative Data Output from ATCC STR Profile Database

| % Match | ATCC Number | Designation | D5S818 | D13S317 | D7S820 | D16S539 | vWA | TH01 | AMEL | TPOX | CSF1PO |
|---|---|---|---|---|---|---|---|---|---|---|---|
| 100 | HTB-26 | MDA-MB-231 Breast Adenocarcinoma Human | 12 | 13 | 8,9 | 12 | 15,18 | 7,9.3 | X | 8,9 | 12,13 |

For alternate database, you may visit https://www.dsmz.de/services/services-human-and-animal-cell-lines/online-str-analysis.html

End of report

**Fragment Analysis**
**- STR Profiling**
**Cell Line Authentication**

Axil Scientific Pte Ltd
2 Tukang Innovation Grove, #06-01, JTC MedTech Hub,
Singapore 618305
T: +65 6775 7318
F: +65 6775 7211
E:

Apical Scientific Sdn Bhd
Lot 7-1 to 7-4, Jalan SP 2/7, Taman Serdang Perdana,
Seksyen 2, 43300 Seri Kembangan, Selangor, Malaysia
T: +603 8943 3252
F: +603 8943 3243
E:

1st BASE
YOUR RESEARCH, OUR BUSINESS
www.base-asia.com

### CUSTOMER INFORMATION

| | | | |
|---|---|---|---|
| Name | Subbulakshmi Karthikeyan | Order ID | 1630 |
| Address | | Date Sample Received | 16-Apr-2026 |
| Email | | Report Date | 23-Apr-2026 |

### METHODOLOGY

Twenty-four short tandem repeat (STR) loci plus the gender determining locus, Amelogenin, were amplified using the commercially available GenePrint® 24 System from Promega. The sample was processed using the Applied Biosystems™ DNA Analyzer. Data were analyzed using GeneMapper® v4.0 software (Applied Biosystems™). Appropriate positive and negative controls were run and confirmed for each sample submitted.

### SAMPLE INFORMATION

| Sample Name | Cell Line Designation |
|---|---|
| 231___WNT11_OE | MDA-MB-231 |

### STR PROFILING RESULTS

| | Test Result for Sample | | | | ATCC Reference Database Profile | |
|---|---|---|---|---|---|---|
| **LOCI** | **231___WNT11_OE** | | | | **MDA-MB-231 Breast Adenocarcinoma Human** | |
| Amelogenin | X | | | | X | |
| D3S1358 | 16 | | | | | |
| D1S1656 | 15 | 17 | | | | |
| D2S441 | 14 | 15 | | | | |
| D10S1248 | 14 | 16 | | | | |
| D13S317 | 13 | | | | 13 | |
| Penta E | 11 | | | | | |
| D16S539 | 12 | | | | 12 | |
| D18S51 | 16 | | | | | |
| D2S1338 | 20 | 21 | | | | |
| CSF1PO | 12 | 13 | | | 12 | 13 |
| Penta D | 11 | 14 | | | | |
| TH01 | 7 | 9.3 | | | 7 | 9.3 |
| vWA | 15 | 18 | | | 15 | 18 |
| D21S11 | 30 | 33.2 | | | | |
| D7S820 | 8 | 9 | | | 8 | 9 |
| D5S818 | 12 | | | | 12 | |
| TPOX | 8 | 9 | | | 8 | 9 |
| DYS391 | | | | | | |
| D8S1179 | 13 | | | | | |
| D12S391 | 17 | 18 | | | | |
| D19S433 | 11 | 14 | | | | |
| FGA | 22 | 23 | | | | |
| D22S1045 | 16 | | | | | |

| | |
|---|---|
| Number of shared alleles between sample and database profile: | 14 |
| Total number of alleles in the database profile: | 14 |
| Percent match between the submitted sample and the database profile: | 100% |

*The allele match algorithm compares the loci highlighted in grey only (8 core loci plus amelogenin).*

### EXPLANATION OF TEST RESULTS

☐ The submitted sample profile is human, but not a match for any profile in the STR database.

☑ The submitted sample profile showed 80% to 100% match for the following ATCC human cell line(s) in the STR database (8 core loci plus Amelogenin): **MDA-MB-231 Breast Adenocarcinoma Human**

☐ The submitted profile is similar to the following ATCC human cell line(s):

☐ The submitted sample is a mixture. Multiple peaks are observed in the STR profiling results.

### ADDITIONAL INFORMATION: Comparative Data Output from ATCC STR Profile Database

| % Match | ATCC Number | Designation | D5S818 | D13S317 | D7S820 | D16S539 | vWA | TH01 | AMEL | TPOX | CSF1PO |
|---|---|---|---|---|---|---|---|---|---|---|---|
| 100 | HTB-26 | MDA-MB-231 Breast Adenocarcinoma Human | 12 | 13 | 8,9 | 12 | 15,18 | 7,9.3 | X | 8,9 | 12,13 |

For alternate database, you may visit https://www.dsmz.de/services/services-human-and-animal-cell-lines/online-str-analysis.html

End of report

**Fragment Analysis**
**- STR Profiling**
**Cell Line Authentication**

Axil Scientific Pte Ltd
2 Tukang Innovation Grove, #06-01, JTC MedTech Hub,
Singapore 618305
T: +65 6775 7318
F: +65 6775 7211
E:

Apical Scientific Sdn Bhd
Lot 7-1 to 7-4, Jalan SP 2/7, Taman Serdang Perdana,
Seksyen 2, 43300 Seri Kembangan, Selangor, Malaysia
T: +603 8943 3252
F: +603 8943 3243
E:

1st BASE
YOUR RESEARCH, OUR BUSINESS
www.base-asia.com

### CUSTOMER INFORMATION

| | | | |
|---|---|---|---|
| Name | Subbulakshmi Karthikeyan | Order ID | 1548 |
| Address | | Date Sample Received | 2-Feb-2026 |
| Email | | Report Date | 5-Feb-2026 |

### METHODOLOGY

Twenty-four short tandem repeat (STR) loci plus the gender determining locus, Amelogenin, were amplified using the commercially available GenePrint® 24 System from Promega. The sample was processed using the Applied Biosystems™ DNA Analyzer. Data were analyzed using GeneMapper® v4.0 software (Applied Biosystems™). Appropriate positive and negative controls were run and confirmed for each sample submitted.

### SAMPLE INFORMATION

| Sample Name | Cell Line Designation |
|---|---|
| Pc3 | Pc3 |

### STR PROFILING RESULTS

| | Test Result for Sample | | | | ExPASy Reference Database Profile | |
|---|---|---|---|---|---|---|
| LOCI | Pc3 | | | | PC-3 | |
| Amelogenin | X | | | | X | |
| D3S1358 | 16 | | | | | |
| D1S1656 | 12 | 16 | | | | |
| D2S441 | 10 | 11 | | | | |
| D10S1248 | 16 | | | | | |
| D13S317 | 11 | | | | 11 | |
| Penta E | 10 | 17 | | | | |
| D16S539 | 11 | | | | 11 | |
| D18S51 | 14 | 15 | | | | |
| D2S1338 | 18 | 20 | | | | |
| CSF1PO | 11 | | | | 11 | |
| Penta D | 9 | | | | | |
| TH01 | 6 | 7 | | | 6 | 7 |
| vWA | 17 | | | | 17 | |
| D21S11 | 29 | 31.2 | | | | |
| D7S820 | 8 | 11 | | | 8 | |
| D5S818 | 13 | | | | 13 | |
| TPOX | 8 | 9 | | | 8 | 9 |
| DYS391 | | | | | | |
| D8S1179 | 13 | | | | | |
| D12S391 | 21 | | | | | |
| D19S433 | 14 | | | | | |
| FGA | 24 | | | | | |
| D22S1045 | 15 | | | | | |

| | |
|---|---|
| Number of shared alleles between sample and database profile: | 11 |
| Total number of alleles in the database profile: | 11 |
| Percent match between the submitted sample and the database profile: | 100% |

*The allele match algorithm compares the loci highlighted in grey only (8 core loci plus amelogenin).*

### EXPLANATION OF TEST RESULTS

[ ] The submitted sample profile is human, but not a match for any profile in the STR database.

[x] The submitted sample profile showed 80% to 100% match for the following ExPASy human cell line(s) in the STR database (8 core loci plus Amelogenin): **PC-3**

[ ] The submitted profile is similar to the following ExPASy human cell line(s):

[ ] The submitted sample is a mixture. Multiple peaks are observed in the STR profiling results.

### ADDITIONAL INFORMATION: Comparative Data Output from ExPASy STR Profile Database

| Accession | Name | Nº Markers | Score | Amel | CSF1PO | D5S818 | D7S820 | D13S317 | D16S539 | TH01 | TPOX | vWA |
|---|---|---|---|---|---|---|---|---|---|---|---|---|
| CVCL_0035 | PC-3 | 9 | 100.00% | X | 11 | 13 | 8 | 11 | 11 | 6,7 | 8,9 | 17 |
| CVCL_XD72 | PC-3-Cas9-576 | 9 | 100.00% | X | 11 | 13 | 8,11 | 11 | 11 | 6,7 | 8,9 | 17 |
| CVCL_XD73 | PC-3-Cas9-577 | 9 | 100.00% | X | 11 | 13 | 8,11 | 11 | 11 | 6,7 | 8,9 | 17 |
| CVCL_XD74 | PC-3-Cas9-578 | 9 | 100.00% | X | 11 | 13 | 8,11 | 11 | 11 | 6,7 | 8,9 | 17 |
| CVCL_XD75 | PC-3-Cas9-579 | 9 | 100.00% | X | 11 | 13 | 8,11 | 11 | 11 | 6,7 | 8,9 | 17 |
| CVCL_J265 | PC-3-Luc | 9 | 100.00% | X | 11 | 13 | 8,11 | 11 | 11 | 6,7 | 8,9 | 17 |
| CVCL_A4BV | PC-3-Luc2 | 9 | 100.00% | X | 11 | 13 | 8,11 | 11 | 11 | 6,7 | 8,9 | 17 |
| CVCL_C169 | PCI-03 | 9 | 100.00% | X | 11 | 13 | 8,11 | 11 | 11 | 7 | 8,9 | 17 |
| CVCL_4778 | PPC-1 | 9 | 100.00% | X | 11 | 13 | 8,11 | 11 | 11 | 6,7 | 8,9 | 17 |
| CVCL_5989 | JHU-019 | 9 | 100.00% | X | 11 | 13 | 8,11 | 11 | 11 | 6,7 | 8,9 | 17 |

For alternate database, you may visit https://www.dsmz.de/services/services-human-and-animal-cell-lines/online-str-analysis.html

End of report

Axil Scientific Pte Ltd
2 Tukang Innovation Grove, #06-01, JTC MedTech Hub,
Singapore 618305
T: +65 6775 7318
F: +65 6775 7211
E:

Apical Scientific Sdn Bhd
Lot 7-1 to 7-4, Jalan SP 2/7, Taman Serdang Perdana,
Seksyen 2, 43300 Seri Kembangan, Selangor, Malaysia
T: +603 8943 3252
F: +603 8943 3243
E:

1st BASE
YOUR RESEARCH, OUR BUSINESS
www.base-asia.com

### CUSTOMER INFORMATION

| | | | |
|---|---|---|---|
| Name | Subbulakshmi Karthikeyan | Order ID | 1630 |
| Address | | Date Sample Received | 16-Apr-2026 |
| Email | | Report Date | 23-Apr-2026 |

### METHODOLOGY

Twenty-four short tandem repeat (STR) loci plus the gender determining locus, Amelogenin, were amplified using the commercially available GenePrint® 24 System from Promega. The sample was processed using the Applied Biosystems™ DNA Analyzer. Data were analyzed using GeneMapper® v4.0 software (Applied Biosystems™). Appropriate positive and negative controls were run and confirmed for each sample submitted.

### SAMPLE INFORMATION

| **Sample Name** | **Cell Line Designation** |
|---|---|
| PC3___EV | PC3 |

### STR PROFILING RESULTS

| | Test Result for Sample | | | | ATCC Reference Database Profile | |
|---|---|---|---|---|---|---|
| **LOCI** | **PC3___EV** | | | | **PC-3Prostate AdenocarcinomaHuman** | |
| Amelogenin | X | | | | X | |
| D3S1358 | 16 | | | | | |
| D1S1656 | 12 | 16 | | | | |
| D2S441 | 10 | 11 | | | | |
| D10S1248 | 16 | | | | | |
| D13S317 | 11 | | | | 11 | |
| Penta E | 10 | 17 | | | | |
| D16S539 | 11 | | | | 11 | |
| D18S51 | 14 | 15 | | | | |
| D2S1338 | 18 | 20 | | | | |
| CSF1PO | 11 | | | | 11 | |
| Penta D | 9 | | | | | |
| TH01 | 6 | 7 | | | 6 | 7 |
| vWA | 17 | | | | 17 | |
| D21S11 | 29 | 31.2 | | | | |
| D7S820 | 8 | 11 | | | 8 | 11 |
| D5S818 | 13 | | | | 13 | |
| TPOX | 8 | 9 | | | 8 | 9 |
| DYS391 | | | | | | |
| D8S1179 | 13 | | | | | |
| D12S391 | 21 | | | | | |
| D19S433 | 14 | | | | | |
| FGA | 24 | | | | | |
| D22S1045 | 15 | | | | | |

| | |
|---|---|
| Number of shared alleles between sample and database profile: | 12 |
| Total number of alleles in the database profile: | 12 |
| Percent match between the submitted sample and the database profile: | 100% |

*The allele match algorithm compares the loci highlighted in grey only (8 core loci plus amelogenin).*

### EXPLANATION OF TEST RESULTS

☐ The submitted sample profile is human, but not a match for any profile in the STR database.

☑ The submitted sample profile showed 80% to 100% match for the following ATCC human cell line(s) in the STR database
(8 core loci plus Amelogenin): **PC-3Prostate AdenocarcinomaHuman**

☐ The submitted profile is similar to the following ATCC human cell line(s):

☐ The submitted sample is a mixture. Multiple peaks are observed in the STR profiling results.

### ADDITIONAL INFORMATION: Comparative Data Output from ATCC STR Profile Database

| % Match | ATCC Number | Designation | D5S818 | D13S317 | D7S820 | D16S539 | vWA | TH01 | AMEL | TPOX | CSF1PO |
|---|---|---|---|---|---|---|---|---|---|---|---|
| 100 | CRL-1435 | PC-3Prostate AdenocarcinomaHuman | 13 | 11 | 8,11 | 11 | 17 | 6,7 | X | 8,9 | 11 |
| 100 | HTB-190 | PPC-1 HUMAN PROSTATE ADENOCARCINOMA | 13 | 11 | 8,11 | 11 | 17 | 6,7 | X | 8,9 | 11 |
| 100 | CRL-1435-LUC2 | PC-3-Luc2; Prostate Adenocarcinoma; Human | 13 | 11 | 8,11 | 11 | 17 | 6,7 | X | 8,9 | 11 |
| 100 | CRL-3470 | PC3-epi Prostate Adenocarcinoma Human | 13 | 11 | 8 | 11 | 17 | 6,7 | X | 8,9 | 11 |
| 100 | CRL-3471 | PC3-emt Prostate Adenocarcinoma Human | 13 | 11 | 8 | 11 | 17 | 6,7 | X | 8,9 | 11 |

For alternate database, you may visit https://www.dsmz.de/services/services-human-and-animal-cell-lines/online-str-analysis.html

End of report

**Fragment Analysis**
**- STR Profiling**
**Cell Line Authentication**

Axil Scientific Pte Ltd
2 Tukang Innovation Grove, #06-01, JTC MedTech Hub,
Singapore 618305
T: +65 6775 7318
F: +65 6775 7211
E:

Apical Scientific Sdn Bhd
Lot 7-1 to 7-4, Jalan SP 2/7, Taman Serdang Perdana,
Seksyen 2, 43300 Seri Kembangan, Selangor, Malaysia
T: +603 8943 3252
F: +603 8943 3243
E:

1st BASE
YOUR RESEARCH, OUR BUSINESS
www.base-asia.com

### CUSTOMER INFORMATION

| | | | |
|---|---|---|---|
| Name | Subbulakshmi Karthikeyan | Order ID | 1630 |
| Address | | Date Sample Received | 16-Apr-2026 |
| Email | | Report Date | 23-Apr-2026 |

### METHODOLOGY

Twenty-four short tandem repeat (STR) loci plus the gender determining locus, Amelogenin, were amplified using the commercially available GenePrint® 24 System from Promega. The sample was processed using the Applied Biosystems™ DNA Analyzer. Data were analyzed using GeneMapper® v4.0 software (Applied Biosystems™). Appropriate positive and negative controls were run and confirmed for each sample submitted.

### SAMPLE INFORMATION

| Sample Name | Cell Line Designation |
|---|---|
| PC3___WNT11_OE | PC3 |

### STR PROFILING RESULTS

| | Test Result for Sample | | | | ATCC Reference Database Profile | |
|---|---|---|---|---|---|---|
| **LOCI** | **PC3___WNT11_OE** | | | | **PC-3Prostate AdenocarcinomaHuman** | |
| Amelogenin | X | | | | X | |
| D3S1358 | 16 | | | | | |
| D1S1656 | 12 | 16 | | | | |
| D2S441 | 10 | 11 | | | | |
| D10S1248 | 16 | | | | | |
| D13S317 | 11 | | | | 11 | |
| Penta E | 10 | 17 | | | | |
| D16S539 | 11 | | | | 11 | |
| D18S51 | 14 | 15 | | | | |
| D2S1338 | 18 | 20 | | | | |
| CSF1PO | 11 | | | | 11 | |
| Penta D | 9 | | | | | |
| TH01 | 6 | 7 | | | 6 | 7 |
| vWA | 17 | | | | 17 | |
| D21S11 | 29 | 31.2 | | | | |
| D7S820 | 8 | 11 | | | 8 | 11 |
| D5S818 | 13 | | | | 13 | |
| TPOX | 8 | 9 | | | 8 | 9 |
| DYS391 | | | | | | |
| D8S1179 | 13 | | | | | |
| D12S391 | 21 | | | | | |
| D19S433 | 14 | | | | | |
| FGA | 24 | | | | | |
| D22S1045 | 15 | | | | | |

| | |
|---|---|
| Number of shared alleles between sample and database profile: | 12 |
| Total number of alleles in the database profile: | 12 |
| Percent match between the submitted sample and the database profile: | 100% |

*The allele match algorithm compares the loci highlighted in grey only (8 core loci plus amelogenin).*

### EXPLANATION OF TEST RESULTS

☐ The submitted sample profile is human, but not a match for any profile in the STR database.

☑ The submitted sample profile showed 80% to 100% match for the following ATCC human cell line(s) in the STR database (8 core loci plus Amelogenin): **PC-3Prostate AdenocarcinomaHuman**

☐ The submitted profile is similar to the following ATCC human cell line(s):

☐ The submitted sample is a mixture. Multiple peaks are observed in the STR profiling results.

### ADDITIONAL INFORMATION: Comparative Data Output from ATCC STR Profile Database

| % Match | ATCC Number | Designation | D5S818 | D13S317 | D7S820 | D16S539 | vWA | TH01 | AMEL | TPOX | CSF1PO |
|---|---|---|---|---|---|---|---|---|---|---|---|
| 100 | CRL-1435 | PC-3Prostate AdenocarcinomaHuman | 13 | 11 | 8,11 | 11 | 17 | 6,7 | X | 8,9 | 11 |
| 100 | HTB-190 | PPC-1 HUMAN PROSTATE ADENOCARCINOMA | 13 | 11 | 8,11 | 11 | 17 | 6,7 | X | 8,9 | 11 |
| 100 | CRL-1435-LUC2 | PC-3-Luc2; Prostate Adenocarcinoma; Human | 13 | 11 | 8,11 | 11 | 17 | 6,7 | X | 8,9 | 11 |
| 100 | CRL-3470 | PC3-epi Prostate Adenocarcinoma Human | 13 | 11 | 8 | 11 | 17 | 6,7 | X | 8,9 | 11 |
| 100 | CRL-3471 | PC3-emt Prostate Adenocarcinoma Human | 13 | 11 | 8 | 11 | 17 | 6,7 | X | 8,9 | 11 |

For alternate database, you may visit https://www.dsmz.de/services/services-human-and-animal-cell-lines/online-str-analysis.html

End of report

**Fragment Analysis**
**- STR Profiling**
**Cell Line Authentication**

Axil Scientific Pte Ltd
2 Tukang Innovation Grove, #06-01, JTC MedTech Hub,
Singapore 618305
T: +65 6775 7318
F: +65 6775 7211
E:

Apical Scientific Sdn Bhd
Lot 7-1 to 7-4, Jalan SP 2/7, Taman Serdang Perdana,
Seksyen 2, 43300 Seri Kembangan, Selangor, Malaysia
T: +603 8943 3252
F: +603 8943 3243
E:

1st BASE
YOUR RESEARCH. OUR BUSINESS
www.base-asia.com

### CUSTOMER INFORMATION

| | | | |
|---|---|---|---|
| Name | Subbulakshmi Karthikeyan | Order ID | 1548 |
| Address | | Date Sample Received | 2-Feb-2026 |
| Email | | Report Date | 5-Feb-2026 |

### METHODOLOGY

Twenty-four short tandem repeat (STR) loci plus the gender determining locus, Amelogenin, were amplified using the commercially available GenePrint® 24 System from Promega. The sample was processed using the Applied Biosystems™ DNA Analyzer. Data were analyzed using GeneMapper® v4.0 software (Applied Biosystems™). Appropriate positive and negative controls were run and confirmed for each sample submitted.

### SAMPLE INFORMATION

| **Sample Name** | **Cell Line Designation** |
|---|---|
| Snb19 | Snb19 |

### STR PROFILING RESULTS

| | **Test Result for Sample** | | | | **ExPASy Reference Database Profile** | |
|---|---|---|---|---|---|---|
| **LOCI** | **Snb19** | | | | **SNB-19** | |
| Amelogenin | X | Y | | | X | |
| D3S1358 | 16 | 17 | | | | |
| D1S1656 | 16 | 17.3 | | | | |
| D2S441 | 11 | | | | | |
| D10S1248 | 14 | | | | | |
| D13S317 | 10 | 11 | | | 10 | 11 |
| Penta E | 7 | 10 | | | | |
| D16S539 | 12 | | | | 12 | |
| D18S51 | 13 | | | | | |
| D2S1338 | 22 | 24 | | | | |
| CSF1PO | 11 | 12 | | | 11 | 12 |
| Penta D | 12 | | | | | |
| TH01 | 9.3 | | | | 9.3 | |
| vWA | 16 | 18 | | | 16 | 18 |
| D21S11 | 29 | | | | | |
| D7S820 | 10 | 12 | | | 10 | |
| D5S818 | 11 | 12 | | | 11 | 12 |
| TPOX | 8 | | | | 8 | |
| DYS391 | 10 | | | | | |
| D8S1179 | 13 | 15 | | | | |
| D12S391 | 17 | 22 | | | | |
| D19S433 | 13 | 15 | | | | |
| FGA | 21 | 25 | | | | |
| D22S1045 | 11 | | | | | |

| | |
|---|---|
| Number of shared alleles between sample and database profile: | 13 |
| Total number of alleles in the database profile: | 13 |
| Percent match between the submitted sample and the database profile: | 100% |

*The allele match algorithm compares the loci highlighted in grey only (8 core loci plus amelogenin).*

### EXPLANATION OF TEST RESULTS

- [ ] The submitted sample profile is human, but not a match for any profile in the STR database.
- [x] The submitted sample profile showed 80% to 100% match for the following ExPASy human cell line(s) in the STR database (8 core loci plus Amelogenin): **SNB-19**
- [ ] The submitted profile is similar to the following ExPASy human cell line(s):
- [ ] The submitted sample is a mixture. Multiple peaks are observed in the STR profiling results.

### ADDITIONAL INFORMATION: Comparative Data Output from ExPASy STR Profile Database

| Accession | Name | Nº Markers | Score | Amel | CSF1PO | D5S818 | D7S820 | D13S317 | D16S539 | TH01 | TPOX | vWA |
|---|---|---|---|---|---|---|---|---|---|---|---|---|
| CVCL_0535 | SNB-19 | 9 | 100.00% | X | 11,12 | 11,12 | 10 | 10,11 | 12 | 9.3 | 8 | 16,18 |
| CVCL_2864 | B2-17 | 9 | 100.00% | X | 11,12 | 11,12 | 10,12 | 10,11 | 12 | 9.3 | 8 | 16,18 |
| CVCL_2800 | KNS-89 | 9 | 100.00% | X | 11,12 | 11,12 | 10,12 | 10,11 | 12 | 9.3 | 8 | 16,18 |
| CVCL_B325 | TK-1 [Human astrocytoma] | 9 | 100.00% | X,Y | 11,12 | 11,12 | 10,12 | 10,11 | 12 | 9.3 | 8 | 16,18 |
| CVCL_0021 | U-251MG | 9 | 100.00% | X | 11,12 | 11 | 10,12 | 10,11 | 12 | 9.3 | 8 | 16,18 |
| CVCL_2809 | U-251MG (KO) | 9 | 100.00% | X | 11,12 | 11 | 10,12 | 10,11 | 12 | 9.3 | 8 | 16,18 |
| CVCL_J269 | U-251MG-Luc | 9 | 100.00% | X,Y | 11,12 | 11,12 | 10,12 | 10,11 | 12 | 9.3 | 8 | 16,18 |
| CVCL_2219 | U-373MG ATCC | 9 | 100.00% | X,Y | 11,12 | 11,12 | 10,12 | 10,11 | 12 | 9.3 | 8 | 16,18 |

For alternate database, you may visit https://www.dsmz.de/services/services-human-and-animal-cell-lines/online-str-analysis.html

End of report

**Fragment Analysis**
**- STR Profiling**
**Cell Line Authentication**

Axil Scientific Pte Ltd
2 Tukang Innovation Grove, #06-01, JTC MedTech Hub,
Singapore 618305
T: +65 6775 7318
F: +65 6775 7211
E:

Apical Scientific Sdn Bhd
Lot 7-1 to 7-4, Jalan SP 2/7, Taman Serdang Perdana,
Seksyen 2, 43300 Seri Kembangan, Selangor, Malaysia
T: +603 8943 3252
F: +603 8943 3243
E:

1st BASE YOUR RESEARCH, OUR BUSINESS
www.base-asia.com

### CUSTOMER INFORMATION

| | | | |
|---|---|---|---|
| Name | Subbulakshmi Karthikeyan | Order ID | 1630 |
| Address | | Date Sample Received | 16-Apr-2026 |
| Email | | Report Date | 23-Apr-2026 |

### METHODOLOGY

Twenty-four short tandem repeat (STR) loci plus the gender determining locus, Amelogenin, were amplified using the commercially available GenePrint® 24 System from Promega. The sample was processed using the Applied Biosystems™ DNA Analyzer. Data were analyzed using GeneMapper® v4.0 software (Applied Biosystems™). Appropriate positive and negative controls were run and confirmed for each sample submitted.

### SAMPLE INFORMATION

| **Sample Name** | **Cell Line Designation** |
|---|---|
| Snb19___EV | Snb19 |

### STR PROFILING RESULTS

| | **Test Result for Sample** | | | | **ATCC Reference Database Profile** | |
|---|---|---|---|---|---|---|
| **LOCI** | **Snb19___EV** | | | | **SNB-19 HUMAN MALIGNANT GLIOBLASTOMA** | |
| Amelogenin | X | Y | | | X | Y |
| D3S1358 | 16 | 17 | | | | |
| D1S1656 | 16 | 17.3 | | | | |
| D2S441 | 11 | | | | | |
| D10S1248 | 14 | | | | | |
| D13S317 | 10 | 11 | | | 10 | 11 |
| Penta E | 7 | 10 | | | | |
| D16S539 | 12 | | | | 12 | |
| D18S51 | 13 | | | | | |
| D2S1338 | 22 | 24 | | | | |
| CSF1PO | 11 | 12 | | | 11 | 12 |
| Penta D | 12 | | | | | |
| TH01 | 9.3 | | | | 9.3 | |
| vWA | 16 | 18 | | | 16 | 18 |
| D21S11 | 29 | | | | | |
| D7S820 | 10 | 12 | | | 10 | 12 |
| D5S818 | 11 | 12 | | | 11 | 12 |
| TPOX | 8 | | | | 8 | |
| DYS391 | 10 | | | | | |
| D8S1179 | 13 | 15 | | | | |
| D12S391 | 17 | 22 | | | | |
| D19S433 | 13 | 15 | | | | |
| FGA | 21 | 25 | | | | |
| D22S1045 | 11 | | | | | |

| | |
|---|---|
| Number of shared alleles between sample and database profile: | 15 |
| Total number of alleles in the database profile: | 15 |
| Percent match between the submitted sample and the database profile: | 100% |

*The allele match algorithm compares the loci highlighted in grey only (8 core loci plus amelogenin).*

### EXPLANATION OF TEST RESULTS

- ☐ The submitted sample profile is human, but not a match for any profile in the STR database.
- ☑ The submitted sample profile showed 80% to 100% match for the following ATCC human cell line(s) in the STR database (8 core loci plus Amelogenin): **SNB-19 HUMAN MALIGNANT GLIOBLASTOMA**
- ☐ The submitted profile is similar to the following ATCC human cell line(s):
- ☐ The submitted sample is a mixture. Multiple peaks are observed in the STR profiling results.

### ADDITIONAL INFORMATION: Comparative Data Output from ATCC STR Profile Database

| % Match | ATCC Number | Designation | D5S818 | D13S317 | D7S820 | D16S539 | vWA | TH01 | AMEL | TPOX | CSF1PO |
|---|---|---|---|---|---|---|---|---|---|---|---|
| 100 | CRL-2219 | SNB-19 HUMAN MALIGNANT GLIOBLASTOMA | 11,12 | 10,11 | 10,12 | 12 | 16,18 | 9.3 | X,Y | 8 | 11,12 |

For alternate database, you may visit https://www.dsmz.de/services/services-human-and-animal-cell-lines/online-str-analysis.html

End of report

**Fragment Analysis**
**- STR Profiling**
**Cell Line Authentication**

Axil Scientific Pte Ltd
2 Tukang Innovation Grove, #06-01, JTC MedTech Hub,
Singapore 618305
T: +65 6775 7318
F: +65 6775 7211
E:

Apical Scientific Sdn Bhd
Lot 7-1 to 7-4, Jalan SP 2/7, Taman Serdang Perdana,
Seksyen 2, 43300 Seri Kembangan, Selangor, Malaysia
T: +603 8943 3252
F: +603 8943 3243
E:

1st BASE
YOUR RESEARCH, OUR BUSINESS
www.base-asia.com

### CUSTOMER INFORMATION

| | | | |
|---|---|---|---|
| Name | Subbulakshmi Karthikeyan | Order ID | 1630 |
| Address | | Date Sample Received | 16-Apr-2026 |
| Email | | Report Date | 23-Apr-2026 |

### METHODOLOGY

Twenty-four short tandem repeat (STR) loci plus the gender determining locus, Amelogenin, were amplified using the commercially available GenePrint® 24 System from Promega. The sample was processed using the Applied Biosystems™ DNA Analyzer. Data were analyzed using GeneMapper® v4.0 software (Applied Biosystems™). Appropriate positive and negative controls were run and confirmed for each sample submitted.

### SAMPLE INFORMATION

| Sample Name | Cell Line Designation |
|---|---|
| Snb19___WNT11_OE | Snb19 |

### STR PROFILING RESULTS

| | Test Result for Sample | | | | ATCC Reference Database Profile | |
|---|---|---|---|---|---|---|
| **LOCI** | **Snb19___WNT11_OE** | | | | **SNB-19 HUMAN MALIGNANT GLIOBLASTOMA** | |
| Amelogenin | X | Y | | | X | Y |
| D3S1358 | 16 | 17 | | | | |
| D1S1656 | 16 | 17.3 | | | | |
| D2S441 | 11 | | | | | |
| D10S1248 | 14 | | | | | |
| D13S317 | 10 | 11 | | | 10 | 11 |
| Penta E | 7 | 10 | | | | |
| D16S539 | 12 | | | | 12 | |
| D18S51 | 13 | | | | | |
| D2S1338 | 22 | 24 | | | | |
| CSF1PO | 11 | 12 | | | 11 | 12 |
| Penta D | 12 | | | | | |
| TH01 | 9.3 | | | | 9.3 | |
| vWA | 16 | 18 | | | 16 | 18 |
| D21S11 | 29 | | | | | |
| D7S820 | 10 | 12 | | | 10 | 12 |
| D5S818 | 11 | 12 | | | 11 | 12 |
| TPOX | 8 | | | | 8 | |
| DYS391 | 10 | | | | | |
| D8S1179 | 13 | 15 | | | | |
| D12S391 | 17 | 22 | | | | |
| D19S433 | 13 | 15 | | | | |
| FGA | 21 | 25 | | | | |
| D22S1045 | 11 | | | | | |

| | |
|---|---|
| Number of shared alleles between sample and database profile: | 15 |
| Total number of alleles in the database profile: | 15 |
| Percent match between the submitted sample and the database profile: | 100% |

*The allele match algorithm compares the loci highlighted in grey only (8 core loci plus amelogenin).*

### EXPLANATION OF TEST RESULTS

☐ The submitted sample profile is human, but not a match for any profile in the STR database.

☑ The submitted sample profile showed 80% to 100% match for the following ATCC human cell line(s) in the STR database (8 core loci plus Amelogenin): **SNB-19 HUMAN MALIGNANT GLIOBLASTOMA**

☐ The submitted profile is similar to the following ATCC human cell line(s):

☐ The submitted sample is a mixture. Multiple peaks are observed in the STR profiling results.

### ADDITIONAL INFORMATION: Comparative Data Output from ATCC STR Profile Database

| % Match | ATCC Number | Designation | D5S818 | D13S317 | D7S820 | D16S539 | vWA | TH01 | AMEL | TPOX | CSF1PO |
|---|---|---|---|---|---|---|---|---|---|---|---|
| 100 | CRL-2219 | SNB-19 HUMAN MALIGNANT GLIOBLASTOMA | 11,12 | 10,11 | 10,12 | 12 | 16,18 | 9.3 | X,Y | 8 | 11,12 |

For alternate database, you may visit https://www.dsmz.de/services/services-human-and-animal-cell-lines/online-str-analysis.html

End of report

**Fragment Analysis**
**- STR Profiling**
**Cell Line Authentication**

Axil Scientific Pte Ltd
2 Tukang Innovation Grove, #06-01, JTC MedTech Hub,
Singapore 618305
T: +65 6775 7318
F: +65 6775 7211
E:

Apical Scientific Sdn Bhd
Lot 7-1 to 7-4, Jalan SP 2/7, Taman Serdang Perdana,
Seksyen 2, 43300 Seri Kembangan, Selangor, Malaysia
T: +603 8943 3252
F: +603 8943 3243
E:

### CUSTOMER INFORMATION

| | | | |
|---|---|---|---|
| Name | Subbulakshmi Karthikeyan | Order ID | 1548 |
| Address | | Date Sample Received | 2-Feb-2026 |
| Email | | Report Date | 5-Feb-2026 |

### METHODOLOGY

Twenty-four short tandem repeat (STR) loci plus the gender determining locus, Amelogenin, were amplified using the commercially available GenePrint® 24 System from Promega. The sample was processed using the Applied Biosystems™ DNA Analyzer. Data were analyzed using GeneMapper® v4.0 software (Applied Biosystems™). Appropriate positive and negative controls were run and confirmed for each sample submitted.

### SAMPLE INFORMATION

| Sample Name | Cell Line Designation |
|---|---|
| MCF7 | MCF7 |

### STR PROFILING RESULTS

| | Test Result for Sample | | | | ExPASy Reference Database Profile | |
|---|---|---|---|---|---|---|
| **LOCI** | **MCF7** | | | | **MCF-7** | |
| Amelogenin | X | | | | X | |
| D3S1358 | 16 | | | | | |
| D1S1656 | 11 | 15.3 | | | | |
| D2S441 | 10 | 14 | | | | |
| D10S1248 | 14 | | | | | |
| D13S317 | 11 | | | | 11 | |
| Penta E | 7 | 12 | | | | |
| D16S539 | 11 | 12 | | | 11 | 12 |
| D18S51 | 14 | | | | | |
| D2S1338 | 21 | 23 | | | | |
| CSF1PO | 10 | | | | 10 | |
| Penta D | 12 | | | | | |
| TH01 | 6 | | | | 6 | |
| vWA | 14 | 15 | | | 14 | 15 |
| D21S11 | 30 | | | | | |
| D7S820 | 8 | 9 | | | 8 | 9 |
| D5S818 | 11 | 12 | | | 11 | 12 |
| TPOX | 9 | 12 | | | 9 | 12 |
| DYS391 | | | | | | |
| D8S1179 | 10 | 14 | | | | |
| D12S391 | 18 | 20 | | | | |
| D19S433 | 13 | 14 | | | | |
| FGA | 23 | 24 | 25 | | | |
| D22S1045 | 15 | 16 | | | | |

| | |
|---|---|
| Number of shared alleles between sample and database profile: | 14 |
| Total number of alleles in the database profile: | 14 |
| Percent match between the submitted sample and the database profile: | 100% |

*The allele match algorithm compares the loci highlighted in grey only (8 core loci plus amelogenin).*

### EXPLANATION OF TEST RESULTS

- [ ] The submitted sample profile is human, but not a match for any profile in the STR database.
- [x] The submitted sample profile showed 80% to 100% match for the following ExPASy human cell line(s) in the STR database (8 core loci plus Amelogenin): **MCF-7**
- [ ] The submitted profile is similar to the following ExPASy human cell line(s):
- [ ] The submitted sample is a mixture. Multiple peaks are observed in the STR profiling results.

### ADDITIONAL INFORMATION: Comparative Data Output from ExPASy STR Profile Database

| Accession | Name | Nº Markers | Score | Amel | CSF1PO | D5S818 | D7S820 | D13S317 | D16S539 | TH01 | TPOX | vWA |
|---|---|---|---|---|---|---|---|---|---|---|---|---|
| CVCL_0031 | MCF-7 | 9 | 100.00% | X | 10 | 11,12 | 8,9 | 11 | 11,12 | 6 | 9,12 | 14,15 |
| CVCL_J259 | MCF-7-Luc | 9 | 100.00% | X | 10 | 11,12 | 8,9 | 11 | 11,12 | 6 | 9,12 | 14,15 |
| CVCL_1D49 | MCF-7/164R-4 | 9 | 100.00% | X | 10 | 11,12 | 8,9 | 11 | 11,12 | 6 | 9,12 | 14,15 |
| CVCL_1D39 | MCF-7/164R-7 | 9 | 100.00% | X | 10 | 11,12 | 8,9 | 11 | 11,12 | 6 | 9,12 | 14,15 |
| CVCL_W536 | MCF-7/182R-6 | 9 | 100.00% | X | 10 | 11,12 | 9 | 11 | 11,12 | 6 | 9,12 | 14,15 |
| CVCL_5A09 | MCF-7/AnaR-1 | 9 | 100.00% | X | 10 | 11,12 | 9 | 11 | 11,12 | 6 | 9,12 | 14,15 |
| CVCL_5A10 | MCF-7/AnaR-2 | 9 | 100.00% | X | 10 | 11,12 | 9 | 11 | 11,12 | 6 | 9,12 | 14,15 |
| CVCL_5A11 | MCF-7/AnaR-3 | 9 | 100.00% | X | 10 | 11,12 | 9 | 11 | 11,12 | 6 | 9,12 | 14,15 |
| CVCL_5A12 | MCF-7/AnaR-4 | 9 | 100.00% | X | 10 | 11,12 | 9 | 11 | 11,12 | 6 | 9,12 | 14,15 |
| CVCL_5A13 | MCF-7/ExeR-1 | 9 | 100.00% | X | 10 | 11,12 | 9 | 11 | 11,12 | 6 | 9,12 | 14,15 |
| CVCL_5A14 | MCF-7/ExeR-2 | 9 | 100.00% | X | 10 | 11,12 | 9 | 11 | 11,12 | 6 | 9,12 | 14,15 |
| CVCL_5A16 | MCF-7/ExeR-4 | 9 | 100.00% | X | 10 | 11,12 | 9 | 11 | 11,12 | 6 | 9,12 | 14,15 |
| CVCL_0U80 | MCF-7/HER2-18 | 9 | 100.00% | X | 10 | 12 | 8,9 | 11 | 11,12 | 6 | 9,12 | 14,15 |
| CVCL_5A17 | MCF-7/LetR-1 | 9 | 100.00% | X | 10 | 11,12 | 9 | 11 | 11,12 | 6 | 9,12 | 14,15 |
| CVCL_5A18 | MCF-7/LetR-2 | 9 | 100.00% | X | 10 | 11,12 | 9 | 11 | 11,12 | 6 | 9,12 | 14,15 |
| CVCL_5A19 | MCF-7/LetR-3 | 9 | 100.00% | X | 10 | 11,12 | 9 | 11 | 11,12 | 6 | 9,12 | 14,15 |
| CVCL_B7P7 | MCF-7/PacR | 9 | 100.00% | X | 10 | 11,12 | 9 | 11 | 11,12 | 6 | 9,12 | 14,15 |
| CVCL_1D47 | MCF-7/S0.5 | 9 | 100.00% | X | 10 | 11,12 | 9 | 11 | 11,12 | 6 | 9,12 | 14,15 |
| CVCL_M436 | MCF-7/TAMR-1 | 9 | 100.00% | X | 10 | 12 | 8,9 | 11 | 11,12 | 6 | 9,12 | 14,15 |
| CVCL_1D42 | MCF-7/TAMR-4 | 9 | 100.00% | X | 10 | 11,12 | 8,9 | 11 | 11,12 | 6 | 9,12 | 14,15 |
| CVCL_1D43 | MCF-7/TAMR-7 | 9 | 100.00% | X | 10 | 11,12 | 8,9 | 11 | 11,12 | 6 | 9,12 | 14,15 |
| CVCL_1D44 | MCF-7/TAMR-8 | 9 | 100.00% | X | 10 | 11,12 | 9 | 11 | 11,12 | 6 | 9,12 | 14,15 |
| CVCL_6860 | MCF-7B | 9 | 100.00% | X | 10 | 11,12 | 8,9 | 11 | 11,12 | 6 | 9,12 | 14,15 |
| CVCL_E2QM | MCF-7GFP | 9 | 100.00% | X | 10 | 12 | 8,9 | 11 | 11,12 | 6 | 9,12 | 14,15 |
| CVCL_1D32 | MCF7 AREc32 | 9 | 100.00% | X | 10 | 11,12 | 9 | 11 | 11,12 | 6 | 9,12 | 14,15 |
| CVCL_A4CH | MCF7 dCas9-KRAB | 9 | 100.00% | X | 10 | 11,12 | 8,9 | 11 | 11,12 | 6 | 9,12 | 14,15 |
| CVCL_XD68 | MCF7-Cas9-542 | 9 | 100.00% | X | 10 | 11,12 | 8,9 | 11 | 11,12 | 6 | 9,12 | 14,15 |
| CVCL_XD69 | MCF7-Cas9-543 | 9 | 100.00% | X | 10 | 11,12 | 8,9 | 11 | 11,12 | 6 | 9,12 | 14,15 |
| CVCL_XD70 | MCF7-Cas9-544 | 9 | 100.00% | X | 10 | 11,12 | 8,9 | 11 | 11,12 | 6 | 9,12 | 14,15 |
| CVCL_A4CI | MCF7-Luc2 [ATCC] | 9 | 100.00% | X | 10 | 11,12 | 8,9 | 11 | 11,12 | 6 | 9,12 | 14,15 |
| CVCL_0413 | MCF7/BUS | 9 | 100.00% | X | 10 | 11,12 | 8,9 | 11 | 11,12 | 6 | 9,12 | 14,15 |
| CVCL_2094 | KPL-1 | 9 | 100.00% | X | 10 | 11,12 | 8,9 | 11 | 11,12 | 6 | 9,12 | 14,15 |

For alternate database, you may visit https://www.dsmz.de/services/services-human-and-animal-cell-lines/online-str-analysis.html

End of report

Axil Scientific Pte Ltd
2 Tukang Innovation Grove, #06-01, JTC MedTech Hub,
Singapore 618305
T: +65 6775 7318
F: +65 6775 7211
E:

Apical Scientific Sdn Bhd
Lot 7-1 to 7-4, Jalan SP 2/7, Taman Serdang Perdana,
Seksyen 2, 43300 Seri Kembangan, Selangor, Malaysia
T: +603 8943 3252
F: +603 8943 3243
E:

1st BASE
YOUR RESEARCH, OUR BUSINESS
www.base-asia.com

### CUSTOMER INFORMATION

| | | | |
|---|---|---|---|
| Name | Subbulakshmi Karthikeyan | Order ID | 1630 |
| Address | | Date Sample Received | 16-Apr-2026 |
| Email | | Report Date | 23-Apr-2026 |

### METHODOLOGY

Twenty-four short tandem repeat (STR) loci plus the gender determining locus, Amelogenin, were amplified using the commercially available GenePrint® 24 System from Promega. The sample was processed using the Applied Biosystems™ DNA Analyzer. Data were analyzed using GeneMapper® v4.0 software (Applied Biosystems™). Appropriate positive and negative controls were run and confirmed for each sample submitted.

### SAMPLE INFORMATION

| **Sample Name** | **Cell Line Designation** |
|---|---|
| MCF7___NT_shRNA | MCF7 |

### STR PROFILING RESULTS

| | **Test Result for Sample** | | | | **ATCC Reference Database Profile** | |
|---|---|---|---|---|---|---|
| **LOCI** | **MCF7___NT_shRNA** | | | | **MCF7 Breast Adenocarcinoma Human** | |
| Amelogenin | X | | | | X | |
| D3S1358 | 16 | | | | | |
| D1S1656 | 11 | 15.3 | | | | |
| D2S441 | 10 | 14 | | | | |
| D10S1248 | 14 | | | | | |
| D13S317 | 11 | | | | 11 | |
| Penta E | 7 | 12 | | | | |
| D16S539 | 11 | 12 | | | 11 | 12 |
| D18S51 | 14 | | | | | |
| D2S1338 | 21 | 23 | | | | |
| CSF1PO | 10 | | | | 10 | |
| Penta D | 12 | | | | | |
| TH01 | 6 | | | | 6 | |
| vWA | 14 | 15 | | | 14 | 15 |
| D21S11 | 30 | | | | | |
| D7S820 | 8 | 9 | | | 8 | 9 |
| D5S818 | 11 | 12 | | | 11 | 12 |
| TPOX | 9 | 12 | | | 9 | 12 |
| DYS391 | | | | | | |
| D8S1179 | 10 | 14 | | | | |
| D12S391 | 18 | 20 | | | | |
| D19S433 | 13 | 14 | | | | |
| FGA | 23 | 24 | 25 | | | |
| D22S1045 | 15 | 16 | | | | |

| | |
|---|---|
| Number of shared alleles between sample and database profile: | 14 |
| Total number of alleles in the database profile: | 14 |
| Percent match between the submitted sample and the database profile: | 100% |

*The allele match algorithm compares the loci highlighted in grey only (8 core loci plus amelogenin).*

### EXPLANATION OF TEST RESULTS

☐ The submitted sample profile is human, but not a match for any profile in the STR database.

☑ The submitted sample profile showed 80% to 100% match for the following ATCC human cell line(s) in the STR database (8 core loci plus Amelogenin): **MCF7 Breast Adenocarcinoma Human**

☐ The submitted profile is similar to the following ATCC human cell line(s):

☐ The submitted sample is a mixture. Multiple peaks are observed in the STR profiling results.

### ADDITIONAL INFORMATION: Comparative Data Output from ATCC STR Profile Database

| % Match | ATCC Number | Designation | D5S818 | D13S317 | D7S820 | D16S539 | vWA | TH01 | AMEL | TPOX | CSF1PO |
|---|---|---|---|---|---|---|---|---|---|---|---|
| 100 | HTB-22 | MCF7 Breast Adenocarcinoma Human | 11,12 | 11 | 8,9 | 11,12 | 14,15 | 6 | X | 9,12 | 10 |
| 100 | BTL-1009 | MCF 7Breast Cancer; Human | 11,12 | 11 | 8,9 | 11,12 | 14,15 | 6 | X | 9,12 | 10 |
| 100 | HTB-22-LUC2 | MCF7-Luc2; Breast Adenocarcinoma; Human | 11,12 | 11 | 8,9 | 11,12 | 14,15 | 6 | X | 9,12 | 10 |
| 100 | HTB-22dCas9-KRAB-MCB | MCF7 dCas9-KRAB; Breast Adenocarcinoma, Human | 11,12 | 11 | 8,9 | 11,12 | 14,15 | 6 | X | 9,12 | 10 |

For alternate database, you may visit https://www.dsmz.de/services/services-human-and-animal-cell-lines/online-str-analysis.html

End of report

**Fragment Analysis**
**- STR Profiling**
**Cell Line Authentication**

Axil Scientific Pte Ltd
2 Tukang Innovation Grove, #06-01, JTC MedTech Hub,
Singapore 618305
T: +65 6775 7318
F: +65 6775 7211
E:

Apical Scientific Sdn Bhd
Lot 7-1 to 7-4, Jalan SP 2/7, Taman Serdang Perdana,
Seksyen 2, 43300 Seri Kembangan, Selangor, Malaysia
T: +603 8943 3252
F: +603 8943 3243
E:

1st BASE
YOUR RESEARCH, OUR BUSINESS
www.base-asia.com

### CUSTOMER INFORMATION

| | | | |
|---|---|---|---|
| Name | Subbulakshmi Karthikeyan | Order ID | 1630 |
| Address | | Date Sample Received | 16-Apr-2026 |
| Email | | Report Date | 23-Apr-2026 |

### METHODOLOGY

Twenty-four short tandem repeat (STR) loci plus the gender determining locus, Amelogenin, were amplified using the commercially available GenePrint® 24 System from Promega. The sample was processed using the Applied Biosystems™ DNA Analyzer. Data were analyzed using GeneMapper® v4.0 software (Applied Biosystems™). Appropriate positive and negative controls were run and confirmed for each sample submitted.

### SAMPLE INFORMATION

| Sample Name | Cell Line Designation |
|---|---|
| MCF7___WNT11_Kd_1 | MCF7 |

### STR PROFILING RESULTS

| | Test Result for Sample | | | | ATCC Reference Database Profile | |
|---|---|---|---|---|---|---|
| **LOCI** | **MCF7___WNT11_Kd_1** | | | | **MCF7 Breast Adenocarcinoma Human** | |
| Amelogenin | X | | | | X | |
| D3S1358 | 16 | | | | | |
| D1S1656 | 11 | 15.3 | | | | |
| D2S441 | 10 | 14 | | | | |
| D10S1248 | 14 | | | | | |
| D13S317 | 11 | | | | 11 | |
| Penta E | 7 | 12 | | | | |
| D16S539 | 11 | 12 | | | 11 | 12 |
| D18S51 | 14 | | | | | |
| D2S1338 | 21 | 23 | | | | |
| CSF1PO | 10 | | | | 10 | |
| Penta D | 12 | | | | | |
| TH01 | 6 | | | | 6 | |
| vWA | 14 | 15 | | | 14 | 15 |
| D21S11 | 30 | | | | | |
| D7S820 | 8 | 9 | | | 8 | 9 |
| D5S818 | 11 | 12 | | | 11 | 12 |
| TPOX | 9 | 12 | | | 9 | 12 |
| DYS391 | | | | | | |
| D8S1179 | 10 | 14 | | | | |
| D12S391 | 18 | 20 | | | | |
| D19S433 | 13 | 14 | | | | |
| FGA | 23 | 24 | 25 | | | |
| D22S1045 | 15 | 16 | | | | |

| | |
|---|---|
| Number of shared alleles between sample and database profile: | 14 |
| Total number of alleles in the database profile: | 14 |
| Percent match between the submitted sample and the database profile: | 100% |

*The allele match algorithm compares the loci highlighted in grey only (8 core loci plus amelogenin).*

### EXPLANATION OF TEST RESULTS

☐ The submitted sample profile is human, but not a match for any profile in the STR database.

☑ The submitted sample profile showed 80% to 100% match for the following ATCC human cell line(s) in the STR database (8 core loci plus Amelogenin): **MCF7 Breast Adenocarcinoma Human**

☐ The submitted profile is similar to the following ATCC human cell line(s):

☐ The submitted sample is a mixture. Multiple peaks are observed in the STR profiling results.

### ADDITIONAL INFORMATION: Comparative Data Output from ATCC STR Profile Database

| % Match | ATCC Number | Designation | D5S818 | D13S317 | D7S820 | D16S539 | vWA | TH01 | AMEL | TPOX | CSF1PO |
|---|---|---|---|---|---|---|---|---|---|---|---|
| 100 | HTB-22 | MCF7 Breast Adenocarcinoma Human | 11,12 | 11 | 8,9 | 11,12 | 14,15 | 6 | X | 9,12 | 10 |
| 100 | BTL-1009 | MCF 7Breast Cancer; Human | 11,12 | 11 | 8,9 | 11,12 | 14,15 | 6 | X | 9,12 | 10 |
| 100 | HTB-22-LUC2 | MCF7-Luc2; Breast Adenocarcinoma; Human | 11,12 | 11 | 8,9 | 11,12 | 14,15 | 6 | X | 9,12 | 10 |
| 100 | HTB-22dCas9-KRAB-MCB | MCF7 dCas9-KRAB; Breast Adenocarcinoma, Human | 11,12 | 11 | 8,9 | 11,12 | 14,15 | 6 | X | 9,12 | 10 |

For alternate database, you may visit https://www.dsmz.de/services/services-human-and-animal-cell-lines/online-str-analysis.html

End of report

**Fragment Analysis**
**- STR Profiling**
**Cell Line Authentication**

Axil Scientific Pte Ltd
2 Tukang Innovation Grove, #06-01, JTC MedTech Hub,
Singapore 618305
T: +65 6775 7318
F: +65 6775 7211
E:

Apical Scientific Sdn Bhd
Lot 7-1 to 7-4, Jalan SP 2/7, Taman Serdang Perdana,
Seksyen 2, 43300 Seri Kembangan, Selangor, Malaysia
T: +603 8943 3252
F: +603 8943 3243
E:

1st BASE
YOUR RESEARCH, OUR BUSINESS
www.base-asia.com

### CUSTOMER INFORMATION

| | | | |
|---|---|---|---|
| Name | Subbulakshmi Karthikeyan | Order ID | 1630 |
| Address | | Date Sample Received | 16-Apr-2026 |
| Email | | Report Date | 23-Apr-2026 |

### METHODOLOGY

Twenty-four short tandem repeat (STR) loci plus the gender determining locus, Amelogenin, were amplified using the commercially available GenePrint® 24 System from Promega. The sample was processed using the Applied Biosystems™ DNA Analyzer. Data were analyzed using GeneMapper® v4.0 software (Applied Biosystems™). Appropriate positive and negative controls were run and confirmed for each sample submitted.

### SAMPLE INFORMATION

| Sample Name | Cell Line Designation |
|---|---|
| MCF7___WNT11_Kd_2 | MCF7 |

### STR PROFILING RESULTS

| | Test Result for Sample | | | | ATCC Reference Database Profile | |
|---|---|---|---|---|---|---|
| **LOCI** | **MCF7___WNT11_Kd_2** | | | | **MCF7 Breast Adenocarcinoma Human** | |
| Amelogenin | X | | | | X | |
| D3S1358 | 16 | | | | | |
| D1S1656 | 11 | 15.3 | | | | |
| D2S441 | 10 | 14 | | | | |
| D10S1248 | 14 | | | | | |
| D13S317 | 11 | | | | 11 | |
| Penta E | 7 | 12 | | | | |
| D16S539 | 11 | 12 | | | 11 | 12 |
| D18S51 | 14 | | | | | |
| D2S1338 | 21 | 23 | | | | |
| CSF1PO | 10 | | | | 10 | |
| Penta D | 12 | | | | | |
| TH01 | 6 | | | | 6 | |
| vWA | 14 | 15 | | | 14 | 15 |
| D21S11 | 30 | | | | | |
| D7S820 | 8 | 9 | | | 8 | 9 |
| D5S818 | 11 | 12 | | | 11 | 12 |
| TPOX | 9 | 12 | | | 9 | 12 |
| DYS391 | | | | | | |
| D8S1179 | 10 | 14 | | | | |
| D12S391 | 18 | 20 | | | | |
| D19S433 | 13 | 14 | | | | |
| FGA | 23 | 24 | 25 | | | |
| D22S1045 | 15 | 16 | | | | |

| | |
|---|---|
| Number of shared alleles between sample and database profile: | 14 |
| Total number of alleles in the database profile: | 14 |
| Percent match between the submitted sample and the database profile: | 100% |

*The allele match algorithm compares the loci highlighted in grey only (8 core loci plus amelogenin).*

### EXPLANATION OF TEST RESULTS

☐ The submitted sample profile is human, but not a match for any profile in the STR database.

☑ The submitted sample profile showed 80% to 100% match for the following ATCC human cell line(s) in the STR database (8 core loci plus Amelogenin): **MCF7 Breast Adenocarcinoma Human**

☐ The submitted profile is similar to the following ATCC human cell line(s):

☐ The submitted sample is a mixture. Multiple peaks are observed in the STR profiling results.

### ADDITIONAL INFORMATION: Comparative Data Output from ATCC STR Profile Database

| % Match | ATCC Number | Designation | D5S818 | D13S317 | D7S820 | D16S539 | vWA | TH01 | AMEL | TPOX | CSF1PO |
|---|---|---|---|---|---|---|---|---|---|---|---|
| 100 | HTB-22 | MCF7 Breast Adenocarcinoma Human | 11,12 | 11 | 8,9 | 11,12 | 14,15 | 6 | X | 9,12 | 10 |
| 100 | BTL-1009 | MCF 7Breast Cancer; Human | 11,12 | 11 | 8,9 | 11,12 | 14,15 | 6 | X | 9,12 | 10 |
| 100 | HTB-22-LUC2 | MCF7-Luc2; Breast Adenocarcinoma; Human | 11,12 | 11 | 8,9 | 11,12 | 14,15 | 6 | X | 9,12 | 10 |
| 100 | HTB-22dCas9-KRAB-MCB | MCF7 dCas9-KRAB; Breast Adenocarcinoma, Human | 11,12 | 11 | 8,9 | 11,12 | 14,15 | 6 | X | 9,12 | 10 |

For alternate database, you may visit https://www.dsmz.de/services/services-human-and-animal-cell-lines/online-str-analysis.html

End of report

Axil Scientific Pte Ltd
2 Tukang Innovation Grove, #06-01, JTC MedTech Hub,
Singapore 618305
T: +65 6775 7318
F: +65 6775 7211
E:

Apical Scientific Sdn Bhd
Lot 7-1 to 7-4, Jalan SP 2/7, Taman Serdang Perdana,
Seksyen 2, 43300 Seri Kembangan, Selangor, Malaysia
T: +603 8943 3252
F: +603 8943 3243
E:

1st BASE
YOUR RESEARCH, OUR BUSINESS
www.base-asia.com

### CUSTOMER INFORMATION

| | | | |
|---|---|---|---|
| Name | Subbulakshmi Karthikeyan | Order ID | 1548 |
| Address | | Date Sample Received | 2-Feb-2026 |
| Email | | Report Date | 5-Feb-2026 |

### METHODOLOGY

Twenty-four short tandem repeat (STR) loci plus the gender determining locus, Amelogenin, were amplified using the commercially available GenePrint® 24 System from Promega. The sample was processed using the Applied Biosystems™ DNA Analyzer. Data were analyzed using GeneMapper® v4.0 software (Applied Biosystems™). Appropriate positive and negative controls were run and confirmed for each sample submitted.

### SAMPLE INFORMATION

| **Sample Name** | **Cell Line Designation** |
|---|---|
| H1299 | H1299 |

### STR PROFILING RESULTS

| | **Test Result for Sample** | | | | **ExPASy Reference Database Profile** | |
|---|---|---|---|---|---|---|
| **LOCI** | **H1299** | | | | **NCI-H1299** | |
| Amelogenin | X | | | | X | |
| D3S1358 | 17 | | | | | |
| D1S1656 | 12 | 15 | | | | |
| D2S441 | 11 | 13 | | | | |
| D10S1248 | 13 | 17 | | | | |
| D13S317 | 12 | | | | 12 | |
| Penta E | 11 | | | | | |
| D16S539 | 12 | 13 | | | 12 | 13 |
| D18S51 | 16 | | | | | |
| D2S1338 | 23 | 24 | | | | |
| CSF1PO | 12 | | | | 12 | |
| Penta D | 13 | | | | | |
| TH01 | 6 | 9.3 | | | 6 | 9.3 |
| vWA | 16 | 18 | | | 16 | 18 |
| D21S11 | 32.2 | | | | | |
| D7S820 | 10 | | | | 10 | |
| D5S818 | 11 | | | | 11 | |
| TPOX | 8 | | | | 8 | |
| DYS391 | | | | | | |
| D8S1179 | 10 | 13 | | | | |
| D12S391 | 21 | 22 | | | | |
| D19S433 | 14 | | | | | |
| FGA | 20 | | | | | |
| D22S1045 | 15 | 16 | | | | |

| | |
|---|---|
| Number of shared alleles between sample and database profile: | 12 |
| Total number of alleles in the database profile: | 12 |
| Percent match between the submitted sample and the database profile: | 100% |

*The allele match algorithm compares the loci highlighted in grey only (8 core loci plus amelogenin).*

### EXPLANATION OF TEST RESULTS

- [ ] The submitted sample profile is human, but not a match for any profile in the STR database.
- [x] The submitted sample profile showed 80% to 100% match for the following ExPASy human cell line(s) in the STR database (8 core loci plus Amelogenin): **NCI-H1299**
- [ ] The submitted profile is similar to the following ExPASy human cell line(s):
- [ ] The submitted sample is a mixture. Multiple peaks are observed in the STR profiling results.

### ADDITIONAL INFORMATION: Comparative Data Output from ExPASy STR Profile Database

| Accession | Name | Nº Markers | Score | Amel | CSF1PO | D5S818 | D7S820 | D13S317 | D16S539 | TH01 | TPOX | vWA |
|---|---|---|---|---|---|---|---|---|---|---|---|---|
| CVCL_0060 | NCI-H1299 | 9 | 100.00% | X | 12 | 11 | 10 | 12 | 12,13 | 6,9.3 | 8 | 16,18 |
| CVCL_XB25 | NCI-H1299-EGFP | 9 | 100.00% | X | 12 | 11 | 10 | 12 | 12,13 | 6,9.3 | 8 | 16,18 |
| CVCL_XE62 | NCI-H1299-Luc2-tdT-2 | 9 | 100.00% | X | 12 | 11 | 10 | 12 | 12,13 | 6,9.3 | 8 | 16,18 |
| CVCL_XB24 | NCI-H1299-mCherry | 9 | 100.00% | X | 12 | 11 | 10 | 12 | 12,13 | 6,9.3 | 8 | 16,18 |
| CVCL_XB26 | NCI-H1299-tdT | 9 | 100.00% | X | 12 | 11 | 10 | 12 | 12,13 | 6,9.3 | 8 | 16,18 |

For alternate database, you may visit https://www.dsmz.de/services/services-human-and-animal-cell-lines/online-str-analysis.html

End of report

**Fragment Analysis**
**- STR Profiling**
**Cell Line Authentication**

Axil Scientific Pte Ltd
2 Tukang Innovation Grove, #06-01, JTC MedTech Hub,
Singapore 618305
T: +65 6775 7318
F: +65 6775 7211
E:

Apical Scientific Sdn Bhd
Lot 7-1 to 7-4, Jalan SP 2/7, Taman Serdang Perdana,
Seksyen 2, 43300 Seri Kembangan, Selangor, Malaysia
T: +603 8943 3252
F: +603 8943 3243
E:

1st BASE
YOUR RESEARCH, OUR BUSINESS
www.base-asia.com

### CUSTOMER INFORMATION

| | | | |
|---|---|---|---|
| Name | Subbulakshmi Karthikeyan | Order ID | 1630 |
| Address | | Date Sample Received | 16-Apr-2026 |
| Email | | Report Date | 23-Apr-2026 |

### METHODOLOGY

Twenty-four short tandem repeat (STR) loci plus the gender determining locus, Amelogenin, were amplified using the commercially available GenePrint® 24 System from Promega. The sample was processed using the Applied Biosystems™ DNA Analyzer. Data were analyzed using GeneMapper® v4.0 software (Applied Biosystems™). Appropriate positive and negative controls were run and confirmed for each sample submitted.

### SAMPLE INFORMATION

| Sample Name | Cell Line Designation |
|---|---|
| H1299___NT_shRNA | H1299 |

### STR PROFILING RESULTS

| | Test Result for Sample | | | | ATCC Reference Database Profile | | |
|---|---|---|---|---|---|---|---|
| **LOCI** | **H1299___NT_shRNA** | | | | **NCI-H1299Lung CarcinomaHuman** | | |
| Amelogenin | X | | | | X | | |
| D3S1358 | 17 | | | | | | |
| D1S1656 | 12 | 15 | | | | | |
| D2S441 | 11 | 13 | | | | | |
| D10S1248 | 13 | 17 | | | | | |
| D13S317 | 12 | | | | 12 | | |
| Penta E | 11 | | | | | | |
| D16S539 | 12 | 13 | 14 | | 12 | 13 | |
| D18S51 | 16 | | | | | | |
| D2S1338 | 23 | 24 | | | | | |
| CSF1PO | 12 | | | | 12 | | |
| Penta D | 13 | | | | | | |
| TH01 | 6 | 9.3 | | | 6 | 9.3 | |
| vWA | 16 | 18 | | | 16 | 17 | 18 |
| D21S11 | 32.2 | | | | | | |
| D7S820 | 10 | | | | 10 | | |
| D5S818 | 11 | | | | 11 | | |
| TPOX | 8 | | | | 8 | | |
| DYS391 | | | | | | | |
| D8S1179 | 10 | 13 | | | | | |
| D12S391 | 21 | 22 | | | | | |
| D19S433 | 14 | | | | | | |
| FGA | 20 | | | | | | |
| D22S1045 | 15 | 16 | | | | | |

| | |
|---|---|
| Number of shared alleles between sample and database profile: | 12 |
| Total number of alleles in the database profile: | 13 |
| Percent match between the submitted sample and the database profile: | 92% |

*The allele match algorithm compares the loci highlighted in grey only (8 core loci plus amelogenin).*

### EXPLANATION OF TEST RESULTS

☐ The submitted sample profile is human, but not a match for any profile in the STR database.

☑ The submitted sample profile showed 80% to 100% match for the following ATCC human cell line(s) in the STR database (8 core loci plus Amelogenin): **NCI-H1299Lung CarcinomaHuman**

☐ The submitted profile is similar to the following ATCC human cell line(s):

☐ The submitted sample is a mixture. Multiple peaks are observed in the STR profiling results.

### ADDITIONAL INFORMATION: Comparative Data Output from ATCC STR Profile Database

| % Match | ATCC Number | Designation | D5S818 | D13S317 | D7S820 | D16S539 | vWA | TH01 | AMEL | TPOX | CSF1PO |
|---|---|---|---|---|---|---|---|---|---|---|---|
| 92 | CRL-5803 | NCI-H1299Lung CarcinomaHuman | 11 | 12 | 10 | 12,13 | 16,17,18 | 6,9.3 | X | 8 | 12 |

For alternate database, you may visit https://www.dsmz.de/services/services-human-and-animal-cell-lines/online-str-analysis.html

End of report

**Fragment Analysis**
**- STR Profiling**
**Cell Line Authentication**

Axil Scientific Pte Ltd
2 Tukang Innovation Grove, #06-01, JTC MedTech Hub,
Singapore 618305
T: +65 6775 7318
F: +65 6775 7211
E:

Apical Scientific Sdn Bhd
Lot 7-1 to 7-4, Jalan SP 2/7, Taman Serdang Perdana,
Seksyen 2, 43300 Seri Kembangan, Selangor, Malaysia
T: +603 8943 3252
F: +603 8943 3243
E:

1st BASE
YOUR RESEARCH, OUR BUSINESS
www.base-asia.com

### CUSTOMER INFORMATION

| | | | |
|---|---|---|---|
| Name | Subbulakshmi Karthikeyan | Order ID | 1630 |
| Address | | Date Sample Received | 16-Apr-2026 |
| Email | | Report Date | 23-Apr-2026 |

### METHODOLOGY

Twenty-four short tandem repeat (STR) loci plus the gender determining locus, Amelogenin, were amplified using the commercially available GenePrint® 24 System from Promega. The sample was processed using the Applied Biosystems™ DNA Analyzer. Data were analyzed using GeneMapper® v4.0 software (Applied Biosystems™). Appropriate positive and negative controls were run and confirmed for each sample submitted.

### SAMPLE INFORMATION

| Sample Name | Cell Line Designation |
|---|---|
| H1299___WNT11_Kd_1 | H1299 |

### STR PROFILING RESULTS

| | Test Result for Sample | | | | ATCC Reference Database Profile | | |
|---|---|---|---|---|---|---|---|
| **LOCI** | **H1299___WNT11_Kd_1** | | | | **NCI-H1299Lung CarcinomaHuman** | | |
| Amelogenin | X | | | | X | | |
| D3S1358 | 17 | | | | | | |
| D1S1656 | 12 | 15 | | | | | |
| D2S441 | 11 | 13 | | | | | |
| D10S1248 | 13 | 17 | | | | | |
| D13S317 | 12 | | | | 12 | | |
| Penta E | 11 | | | | | | |
| D16S539 | 12 | 13 | 14 | | 12 | 13 | |
| D18S51 | 16 | | | | | | |
| D2S1338 | 23 | 24 | | | | | |
| CSF1PO | 12 | | | | 12 | | |
| Penta D | 13 | | | | | | |
| TH01 | 6 | 9.3 | | | 6 | 9.3 | |
| vWA | 16 | 18 | | | 16 | 17 | 18 |
| D21S11 | 32.2 | | | | | | |
| D7S820 | 10 | | | | 10 | | |
| D5S818 | 11 | | | | 11 | | |
| TPOX | 8 | | | | 8 | | |
| DYS391 | | | | | | | |
| D8S1179 | 10 | 13 | | | | | |
| D12S391 | 21 | 22 | | | | | |
| D19S433 | 14 | | | | | | |
| FGA | 20 | | | | | | |
| D22S1045 | 15 | 16 | | | | | |

| | |
|---|---|
| Number of shared alleles between sample and database profile: | 12 |
| Total number of alleles in the database profile: | 13 |
| Percent match between the submitted sample and the database profile: | 92% |

*The allele match algorithm compares the loci highlighted in grey only (8 core loci plus amelogenin).*

### EXPLANATION OF TEST RESULTS

☐ The submitted sample profile is human, but not a match for any profile in the STR database.

☑ The submitted sample profile showed 80% to 100% match for the following ATCC human cell line(s) in the STR database
(8 core loci plus Amelogenin): **NCI-H1299Lung CarcinomaHuman**

☐ The submitted profile is similar to the following ATCC human cell line(s):

☐ The submitted sample is a mixture. Multiple peaks are observed in the STR profiling results.

### ADDITIONAL INFORMATION: Comparative Data Output from ATCC STR Profile Database

| % Match | ATCC Number | Designation | D5S818 | D13S317 | D7S820 | D16S539 | vWA | TH01 | AMEL | TPOX | CSF1PO |
|---|---|---|---|---|---|---|---|---|---|---|---|
| 92 | CRL-5803 | NCI-H1299Lung CarcinomaHuman | 11 | 12 | 10 | 12,13 | 16,17,18 | 6,9.3 | X | 8 | 12 |

For alternate database, you may visit https://www.dsmz.de/services/services-human-and-animal-cell-lines/online-str-analysis.html

End of report

**Fragment Analysis**
**- STR Profiling**
**Cell Line Authentication**

Axil Scientific Pte Ltd
2 Tukang Innovation Grove, #06-01, JTC MedTech Hub,
Singapore 618305
T: +65 6775 7318
F: +65 6775 7211
E:

Apical Scientific Sdn Bhd
Lot 7-1 to 7-4, Jalan SP 2/7, Taman Serdang Perdana,
Seksyen 2, 43300 Seri Kembangan, Selangor, Malaysia
T: +603 8943 3252
F: +603 8943 3243
E:

www.base-asia.com

### CUSTOMER INFORMATION

| | | | |
|---|---|---|---|
| Name | Subbulakshmi Karthikeyan | Order ID | 1630 |
| Address | | Date Sample Received | 16-Apr-2026 |
| Email | | Report Date | 23-Apr-2026 |

### METHODOLOGY

Twenty-four short tandem repeat (STR) loci plus the gender determining locus, Amelogenin, were amplified using the commercially available GenePrint® 24 System from Promega. The sample was processed using the Applied Biosystems™ DNA Analyzer. Data were analyzed using GeneMapper® v4.0 software (Applied Biosystems™). Appropriate positive and negative controls were run and confirmed for each sample submitted.

### SAMPLE INFORMATION

| **Sample Name** | **Cell Line Designation** |
|---|---|
| H1299___WNT11_Kd_2 | H1299 |

### STR PROFILING RESULTS

| | **Test Result for Sample** | | | | **ATCC Reference Database Profile** | | |
|---|---|---|---|---|---|---|---|
| **LOCI** | **H1299___WNT11_Kd_2** | | | | **NCI-H1299Lung CarcinomaHuman** | | |
| Amelogenin | X | | | | X | | |
| D3S1358 | 17 | | | | | | |
| D1S1656 | 12 | 15 | 16.2 | | | | |
| D2S441 | 11 | 13 | | | | | |
| D10S1248 | 13 | 17 | | | | | |
| D13S317 | 12 | | | | 12 | | |
| Penta E | 11 | | | | | | |
| D16S539 | 12 | 13 | | | 12 | 13 | |
| D18S51 | 16 | | | | | | |
| D2S1338 | 23 | 24 | | | | | |
| CSF1PO | 12 | | | | 12 | | |
| Penta D | 13 | | | | | | |
| TH01 | 6 | 10 | | | 6 | 9.3 | |
| vWA | 16 | 18 | | | 16 | 17 | 18 |
| D21S11 | 32.2 | | | | | | |
| D7S820 | 10 | | | | 10 | | |
| D5S818 | 11 | | | | 11 | | |
| TPOX | 8 | | | | 8 | | |
| DYS391 | | | | | | | |
| D8S1179 | 10 | 13 | | | | | |
| D12S391 | 21 | 22 | | | | | |
| D19S433 | 14 | | | | | | |
| FGA | 20 | | | | | | |
| D22S1045 | 15 | 16 | | | | | |

| | |
|---|---|
| Number of shared alleles between sample and database profile: | 11 |
| Total number of alleles in the database profile: | 13 |
| Percent match between the submitted sample and the database profile: | 85% |

*The allele match algorithm compares the loci highlighted in grey only (8 core loci plus amelogenin).*

### EXPLANATION OF TEST RESULTS

☐ The submitted sample profile is human, but not a match for any profile in the STR database.

☑ The submitted sample profile showed 80% to 100% match for the following ATCC human cell line(s) in the STR database (8 core loci plus Amelogenin): **NCI-H1299Lung CarcinomaHuman**

☐ The submitted profile is similar to the following ATCC human cell line(s):

☐ The submitted sample is a mixture. Multiple peaks are observed in the STR profiling results.

### ADDITIONAL INFORMATION: Comparative Data Output from ATCC STR Profile Database

| % Match | ATCC Number | Designation | D5S818 | D13S317 | D7S820 | D16S539 | vWA | TH01 | AMEL | TPOX | CSF1PO |
|---|---|---|---|---|---|---|---|---|---|---|---|
| 85 | CRL-5803 | NCI-H1299Lung CarcinomaHuman | 11 | 12 | 10 | 12,13 | 16,17,18 | 6,9.3 | X | 8 | 12 |

For alternate database, you may visit https://www.dsmz.de/services/services-human-and-animal-cell-lines/online-str-analysis.html

End of report
